# Body-Axis Reorientation in Regenerating *Hydra* under Geometric Confinement

**DOI:** 10.64898/2026.06.25.734673

**Authors:** Ariel Westfried, Liora Garion, Marko Popović, Kinneret Keren

## Abstract

Defining a body axis is a central aspect of animal morphogenesis. During regeneration from excised *Hydra* tissue pieces, the newly formed body axis typically preserves the orientation of the parent body axis and aligns with the inherited nematic organization of the supracellular actomyosin fibers. Here we show that this inherited orientation can be overridden by geometric confinement. Tissue spheroids confined in narrow cylindrical channels in a frustrating configuration, with the inherited axis initially perpendicular to the channel, regenerate with their body axis aligned along the channel. Using high-resolution live imaging we show that this reorientation is accompanied by remodeling of the nematic fiber organization. New fibers form parallel to the channel axis in the initially disordered closure regions, creating sharp domain boundaries with the inherited transverse fibers. These domain boundaries subsequently propagate, with perpendicular fibers dissolving and new fibers forming along the channel axis. The confined tissue behaves as a solid-like active nematic material, storing anisotropic strain over long timescales while allowing nematic reorganization relative to the material frame. Our results suggest that coupling between tissue strain and nematic alignment contributes to fiber reorientation and body-axis patterning, highlighting how external mechanical constraints can redirect the body axis during morphogenesis.

---

The establishment of a body axis is a fundamental step in animal morphogenesis, providing the basic framework that guides tissue patterning and body-plan formation. This process is central during both embryonic development and whole-body regeneration in organisms such as *Hydra*. In many systems, axis formation is guided by pre-existing directional cues, such as maternal determinants in the oocyte, inherited polarity in the tissue, or external signals. In other cases, axis formation involves *de novo* symmetry breaking, in which an initially symmetric system develops polarity through self-organized interactions^1^. Although biochemical signaling pathways play a central role in axis formation, increasing evidence shows that mechanical cues can also influence this process^2,3^. For example, in early mammalian embryos, cell contractility, cell surface tension, and mechanical interactions between cells have been implicated in the emergence and refinement of embryonic axes^4^.

The actomyosin cytoskeleton is the main force generator in multicellular animal systems. Parallel arrays of contractile actomyosin fibers are a common organizational theme seen across a variety of animals, including *Hydra*, worms, insects and vertebrate species^5–10^. Multiple fibers arrange into nematic supracellular arrays by linking actomyosin bundles between neighboring cells. These supracellular nematic fiber arrays are not merely structural motifs but play central roles in shaping tissue-scale morphology and dynamics^6,10^. Coordinated motor activity within these arrays induces synchronized contractions, linking cellular-level contractility to tissue-scale force generation. Such tissue-spanning contractions drive processes ranging from cardiac beating^8^ and gut peristalsis in vertebrates^7^ to the shaping of the oocyte within the *Drosophila* egg chamber^5^. The organizational principles that govern the formation, alignment, and maintenance of these contractile actomyosin fiber arrays are still not well understood, and likely involve extensive feedback between fiber organization, force generation and morphology. For example, in the developing gut, differential growth generates residual strains that have been shown to orient the inner smooth muscle layer circumferentially, and contractions of this inner layer subsequently provide mechanical cues that align the outer smooth muscle layer longitudinally^7^.

Here, we study *Hydra*, famous for its extraordinary regenerative capabilities, to investigate how geometrical confinement influences the development of the body plan during regeneration, focusing on the determination of the body axis. *Hydra* has a simple uniaxial body plan with a bilayered epithelial body wall. Both epithelial layers contain supracellular contractile actomyosin fibers organized in nematic arrays that span the entire organism^9,11,12^. The fibers in the outer ectoderm layer are aligned with the body axis, whereas fibers in the inner endoderm layer form circumferential rings. Individual cells are ∼20 μm in diameter, whereas the typical spacing between fibers is smaller (∼3.5 μm for ectodermal fibers^13,14^), so each cell contains several fibers. Small excised tissue pieces or aggregates of dissociated cells can regenerate into complete animals within days^15^. During this whole-body regeneration, contractile fiber arrays develop, either from partially inherited structures in excised tissue pieces^9,11,12,16^ or completely *de novo* in regenerating cell aggregates^14^. The alignment of the supracellular actomyosin fibers in excised tissue pieces confers a structural memory that persists during regeneration and aligns with the body axis in the new animal^11,16^. Moreover, topological defects in the nematic alignment of the actomyosin fibers emerge early during regeneration and coincide with the formation sites of morphological features in the regenerated animal^9,12^.

A central step in axial patterning during *Hydra* regeneration is the emergence of a head organizer, located at the tip of the mouth in mature animals^17^. The establishment of a head organizer has largely been attributed to biochemical signaling associated with the Wnt/β-catenin pathway and its autoregulatory dynamics^18–20^. Notably, the Wnt/β-catenin signaling center at the head organizer coincides with an aster-shaped, +1 topological defect in the actomyosin fiber alignment^12^. Functionally, both features are essential: specific biochemical signals are needed to direct the differentiation of specialized cells in the head region, while the aster-shaped +1 defect enables mouth opening upon fiber contraction^21^. However, despite their functional importance and striking spatial correlation, it remains unclear how actomyosin fiber organization and biochemical signals are coupled, and how this interplay guides axis specification.

Confinement can be used to restrict and direct tissue dynamics along an externally-specified axis. Previous work from our lab has shown that axial patterning in regenerating *Hydra* can be influenced by confinement within narrow cylindrical channels^22^. Specifically, we showed that confinement of tissue spheroids promoted the regeneration of *Hydra* with a multi-axial morphology. Similar results were obtained by confining bisected *Hydra* between two planar surfaces^23^. Together, these studies show that external geometric constraints can strongly perturb the regenerated body plan, motivating us to ask how confinement affects the nematic fiber organization and how this reorganization relates to the establishment and alignment of the regenerated body axis.

Here, we confine regenerating *Hydra* spheroids in narrow cylindrical channels in a frustrated configuration, where the channel axis is perpendicular to the inherited body axis of the parent animal^22^. We show that in this perpendicular configuration, both the primary fiber orientation and the regenerated body axis reorient relative to the tissue frame of reference and become aligned with the channel axis. We characterize tissue strain patterns and fiber reorientation dynamics, linking cellular-level organization to tissue-scale dynamics. We find that confinement generates highly anisotropic elastic strain in the tissue that persists for days. From a physical perspective, our results show that a regenerating *Hydra* behave as an active nematic solid, in which the nematic actomyosin fibers reorganize within the tissue frame of reference and become aligned with the channel axis and the primary strain direction. From a biological perspective, we show that we can redirect the regenerated body axis along an externally imposed axis defined by the channel, transverse to the inherited axis from the parent animal.

## Body axis reorientation in confined regenerating Hydra under frustration

To study how geometric confinement influences body-axis formation, we introduce regenerating *Hydra* tissue spheroids into narrow cylindrical channels in a frustrating configuration, with their original body axis oriented perpendicular to the “easy axis” defined by the channel geometry^22^ (Fig. 1A). Excised tissue rings fold into sealed hollow spheroids, with roughly radial symmetry around the original body axis. These spheroids contain an inherited nematic array of supracellular ectodermal fibers aligned with the parent animal’s body axis, as well as two closure regions on their originally head-facing and foot-facing sides. These closure regions are largely devoid of aligned supracellular ectodermal fibers at the time of channel entry. The folded spheroids are then inserted into narrow channels (180-300 μm in diameter; see Methods), 3-8 hours after excision. For the size range of tissue rings used here, the sealed spheroids are typically oblate, so flow tends to align them during insertion with their original body axis transverse to the channel axis (Fig. 1A).

**Figure 1.**
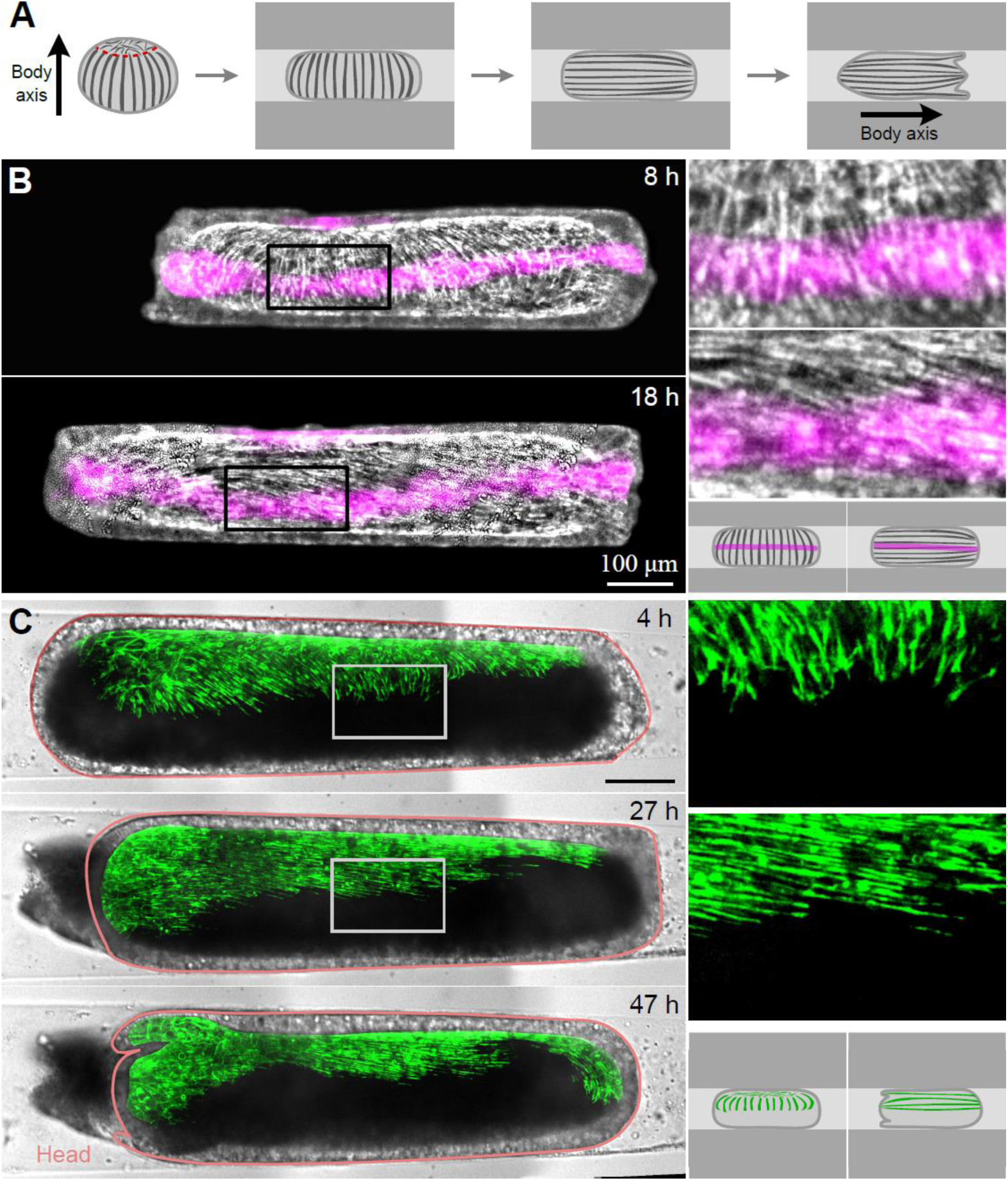
Axis reorientation in regenerating Hydra confined in a frustrating perpendicular configuration. (A) Schematic illustration of an oblate tissue spheroid generated by folding of an excised tissue ring. The folded spheroid has two closure regions at the top (dashed red line) and bottom (not visible). The spheroid is then inserted into a narrow cylindrical channel, where it is lodged with its original body axis transverse to the channel axis. In this side view of the confined spheroid, the two closure regions are not visible. The spheroid subsequently regenerates with its body axis forming along the channel axis. (B) Axis reorientation in a regenerating Hydra labeled with a photoactivated fluorescent stripe. Left: Images of Lifeact-GFP (gray) overlaid with a photoactivated tissue marker (magenta; Abberior CAGE 552) from a time-lapse movie of a regenerating Hydra confined in a perpendicular configuration 8 hours and 18 hours after excision (Movie 2). Right top: Zoomed images of the boxed regions on the left images. Right bottom: Schematic illustration of a confined tissue spheroid labeled with a fluorescent stripe reorienting its axis. The actomyosin fibers are perpendicular to the labeled stripe at the early time point and parallel to it at the later one, while the stripe maintains its orientation relative to the channel, indicating reorientation of the fibers within the tissue. (C) Reorientation of the actomyosin fibers and the body axis in a chimerically-labeled regenerating Hydra. Left: Brightfield and fluorescence overlaid images of a half-labeled tissue spheroid confined in a perpendicular configuration, showing the fiber organization at early (top), intermediate (middle) and late (bottom) time points in the regeneration process (Movie 3). The inherited actomyosin fibers are initially transverse to the graft seam and the channel axis (top), reorient along the channel axis (middle), and eventually the animal regenerates along this axis (bottom). Right top: Zoomed fluorescence images of the boxed regions on the left images. Right bottom: Schematic illustration of a confined chimeric tissue spheroid reorienting its axis.

Once inserted, the spheroids are confined within the channels and deform into an elongated shape, with a cylindrical central region pressed against the channel walls and rounded caps facing the two open ends of the channel (Fig. 1A). Since the tissue does not adhere to the channel walls, the forces exerted by the walls are expected to be predominantly normal to the tissue surface. Tangential forces can, however, arise from frictional contact with the walls. The confined samples regenerate into mature, functional animals within 40-70 hours after excision, exhibiting cycles of slow osmotic inflations and abrupt ruptures (Fig. S1; Movie 1), as seen in unconfined tissues^24,25^. This behavior produces a saw-tooth pattern in the spheroids’ projected area over time (Fig. S1C). To assess the regenerated body plan, samples are removed from the channel 3 days after excision and the regenerated animals are examined outside the confining environment (Fig. S2; Methods).

To follow body-axis formation and relate it to the orientation of the parent animal’s body axis, we use transgenic *Hydra* expressing Lifeact-GFP in the ectoderm to visualize the supracellular ectodermal fibers, paired with a photoactivated marker that labels a stripe of tissue initially parallel to the channel axis (Fig. 1B; Methods). In most confined samples, the tissue does not appreciably rotate within the channel during the first 24 hours after excision. As such, the strain imposed by the channel maintains a well-defined orientation in the tissue frame of reference. Moreover, the approximate alignment between the laboratory and tissue frames of reference simplifies tracking of nematic reorganization over time. In the perpendicular configuration, the inherited fiber array in the central part of the confined spheroid is initially oriented transverse to the channel axis but later becomes predominantly aligned with the channel axis (Fig. 1B, Movie 2). Overall, for samples confined in the perpendicular configuration with an aspect ratio (sample length over channel diameter) larger than 2 (N=31), 28 out of 31 did not rotate appreciably within the channel during the first 24 hours after excision. In all these samples (28/28), tissue regions that initially contained inherited fibers transverse to the channel axis developed fiber arrays aligned with the channel axis, accompanied by reorientation of the regenerated body axis (Fig. 1B).

As a complementary approach, we use axially labeled chimeric animals in which only part of the tissue expresses Lifeact-GFP in the ectoderm (Figs. 1C, S3; Movies 3,4). These animals are generated by grafting bisected *Hydra* that are differentially labeled, such that the graft seam is transverse to the parent animal’s body axis and to the primary nematic orientation of the supracellular fibers (Fig. S3). A tissue ring excised around the graft seam region of a chimeric animal folds into a half-labeled spheroid (Fig. S3C). In these spheroids, the graft seam provides a persistent spatial landmark for the original tissue orientation, allowing us to determine that confined spheroids do not appreciably rotate while the nematic fiber orientation becomes aligned with the channel axis (Fig. 1C). Thus, fibers initially oriented primarily transverse to the graft seam reorient to become parallel to it. At later stages of regeneration, we often observe extensive rotations within the channel (Fig. S4; Movie 5), which correlate with elongation of the tissue along the regenerated body axis^26,27^. The axial elongation of the regenerating sample is accompanied by a reduction in tissue diameter, which decreases tissue contact with the channel walls. This reduces frictional constraints and allows the tissue to rotate more freely within the channel. Overall, our observations demonstrate that although excised tissue rings typically inherit their primary fiber orientation and body-axis orientation from the parent animal, under frustrating confinement *Hydra* tissues can reorient both the fiber orientation and their body axis within the tissue frame of reference (Fig. 1B,C).

## Actomyosin fibers dynamics during axis reorientation

To follow how the nematic actomyosin fibers reorient under frustration, we use high-resolution live imaging of confined *Hydra* tissues expressing Lifeact-GFP in the ectoderm (Fig. 2; Movie 6). This allows us to visualize the dynamics of supracellular fibers on the basal side of the ectoderm together with apical junctions delineating cells at the apical surface (Fig. 2D). By manually tracking individual cells, we establish a tissue frame of reference that allows us to resolve fiber reorganization dynamics. In the frustrated perpendicular configuration, the two closure regions, which are initially disordered, are pressed against the channel wall on opposite sides of the confined tissue. The tissue between these closure regions contains the inherited array of nematic fibers aligned with the original body axis. In the central part of the confined tissue, these inherited fibers are aligned transverse to the channel axis, whereas at the far ends of the tissue, the fibers curve toward the nearby closure region (Fig. 2A,F). The spheroid’s original symmetry axis can be oriented at different angles relative to the imaging plane (Fig. S5). Depending on this angle, the imaged view may show primarily the inherited fibers (Fig. S5B), a nearly head-on view of the closure region (Fig. S5D), or, most commonly, an intermediate view in which both inherited fibers and one of the closure regions are visible (Figs. 2, S5C).

**Figure 2.**
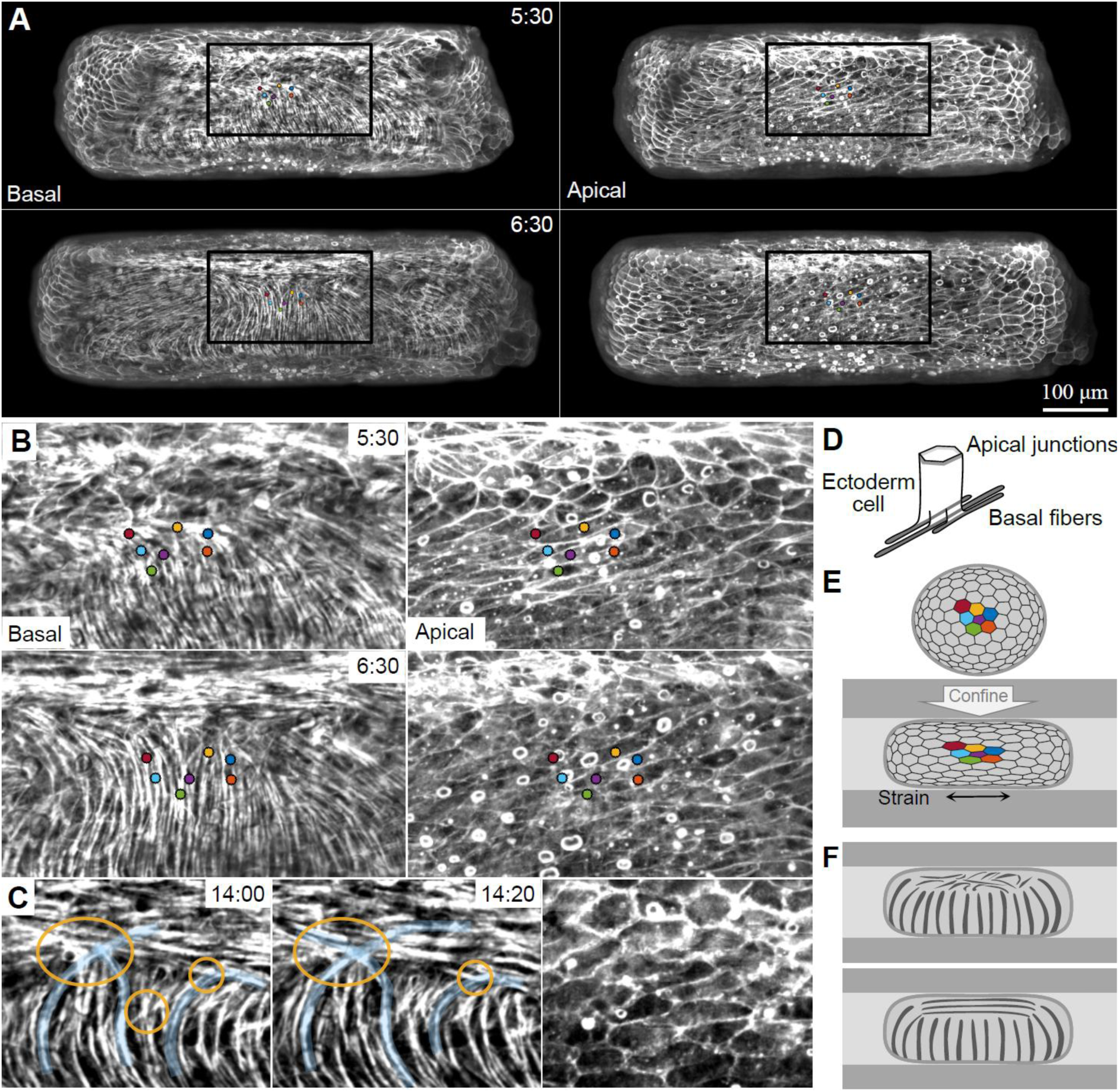
Newly formed fibers in closure regions align with the channel axis. (A) Spinning-disk confocal projected images of the ectodermal basal and apical surfaces in a tissue spheroid expressing Lifeact-GFP confined in a perpendicular configuration at 5.5 and 6.5 hours past excision (∼2.5 and ∼3.5 hours after insertion to channel, respectively; Movie 6). (B) Zoomed images of the boxed region in (A). Colored dots depict individual cells that were manually tracked over time. (C) Fiber organization at the domain boundary. Left: zoomed images of the basal surface at the domain boundary at consecutive time points. Teal highlights actomyosin fibers that gradually change their orientation, and orange circles depict regions with criss-crossing fibers. Right: image of the apical surface in the same region. (D) Schematic of an ectodermal cell. Apical junctions delineate the cell boundary at the apical surface, and aligned actomyosin fibers span the basal surface. (E) Schematic illustrations of the apical surface of a tissue spheroid before (top) and after (bottom) confinement. Cell shape anisotropy reflects the tissue strain induced by confinement. (F) Schematic illustrations of the basal surface of a confined tissue spheroid showing the closure region at an early time point where it is still disordered (top), and a later time point after the formation of a parallel fiber array aligned with the channel axis (bottom).

The closure regions in confined spheroids are initially disordered. We observe reformation of aligned fibers in these regions, with an orientation parallel to the channel axis, in all samples examined (Fig. 2A,B; N=35). This process occurs relatively fast, with an ordered parallel domain appearing within 2-5 hours after confinement. As a result, two “island” domains of parallel fibers that are aligned with the channel axis form at the antipodal closure regions, while the surrounding inherited fibers remain relatively stable over this time frame. Within the cylindrical section of the sample, the boundaries between parallel and perpendicular domains extend along the channel axis (Fig. 2B,F). Towards the sides of the sample, as the parallel fibers form in the closure regions, they connect to the inherited fiber ends near the two caps facing the open ends of the channel.

The boundary between the parallel and perpendicular domains is abrupt on the tissue scale (Fig. 2C). The fiber organization at the domain boundary exhibits two distinct local patterns. First, we observe individual supracellular fibers that connect both domains by changing their orientation in a continuous manner (teal highlights in Fig. 2C). These fibers can bend in either direction as they reorient, turning clockwise or counterclockwise, while maintaining continuity across the interface. Second, we observe crossing fibers (orange circles in Fig. 2C). That is, within the same local region, we find fibers with roughly perpendicular orientations, forming a criss-cross pattern. The fibers undergo recurring contractions and small local rearrangements on the time scale of minutes. Over longer time scales, these dynamics lead to movement of the domain boundary and changes in the local fiber structure and orientation. Notably, although the boundary moves and the fibers rearrange, the transition between the two domains remains abrupt on the tissue scale.

The confined tissue is subjected primarily to normal forces from the channel walls. Because the tissue does not adhere to the channel walls, the only tangential forces it experiences arise from friction. We can infer the primary strain orientation within the tissue from the cell shapes, which are visible in images of the apical surface. Shortly after insertion, cells exhibit pronounced anisotropy: they are elongated along the channel axis and compressed circumferentially (Figs. 2B, S6). This deformation pattern is consistent with an elastic cellular shell forced into a cylindrical geometry, where cells maintain their neighbors, and hence the number of cells around the circumference and along the axial direction should not change upon confinement (Fig. 2E). Instead, the imposed geometrical deformation generates anisotropic cellular strain, with extension along the channel axis and compression in the circumferential direction (Fig. 2E), as observed (Fig. 2B).

The newly formed fibers in the closure region are aligned along the channel axis, which is also the principal direction of stress and strain imposed by confinement (Fig. 2B). This raises the possibility that fiber formation is guided by the anisotropic stress or strain field in the tissue, with fibers assembling preferentially along its principal axis. Notably, the strain anisotropy inferred from the cell shapes persists over timescales much longer than those required for fiber formation: while fibers form within the first 2-5 hours after confinement, the anisotropic cell shapes persist much longer (Fig. S6). Moreover, even after 3 days in the channels, tissue samples immediately recoil and become rounder upon release from confinement (Fig. S7).

To follow the domain boundary dynamics, we color-code the local orientation angle (Figs. 3,4; Movie 6). This color-coding enhances the contrast between neighboring domains and highlights the domain boundary (Figs. 3D, 4A,C). We follow the reorganization of basal fibers over time, using tracked cells in the apical surface to define a tissue frame of reference. This analysis reveals that domain-boundary propagation results from nematic rearrangements within the tissue, rather than from bulk tissue motion. The propagation of the domain boundary occurs normal to the boundary, resulting in the displacement of the domain boundary in a direction perpendicular to the channel axis (Fig. 3).

**Figure 3.**
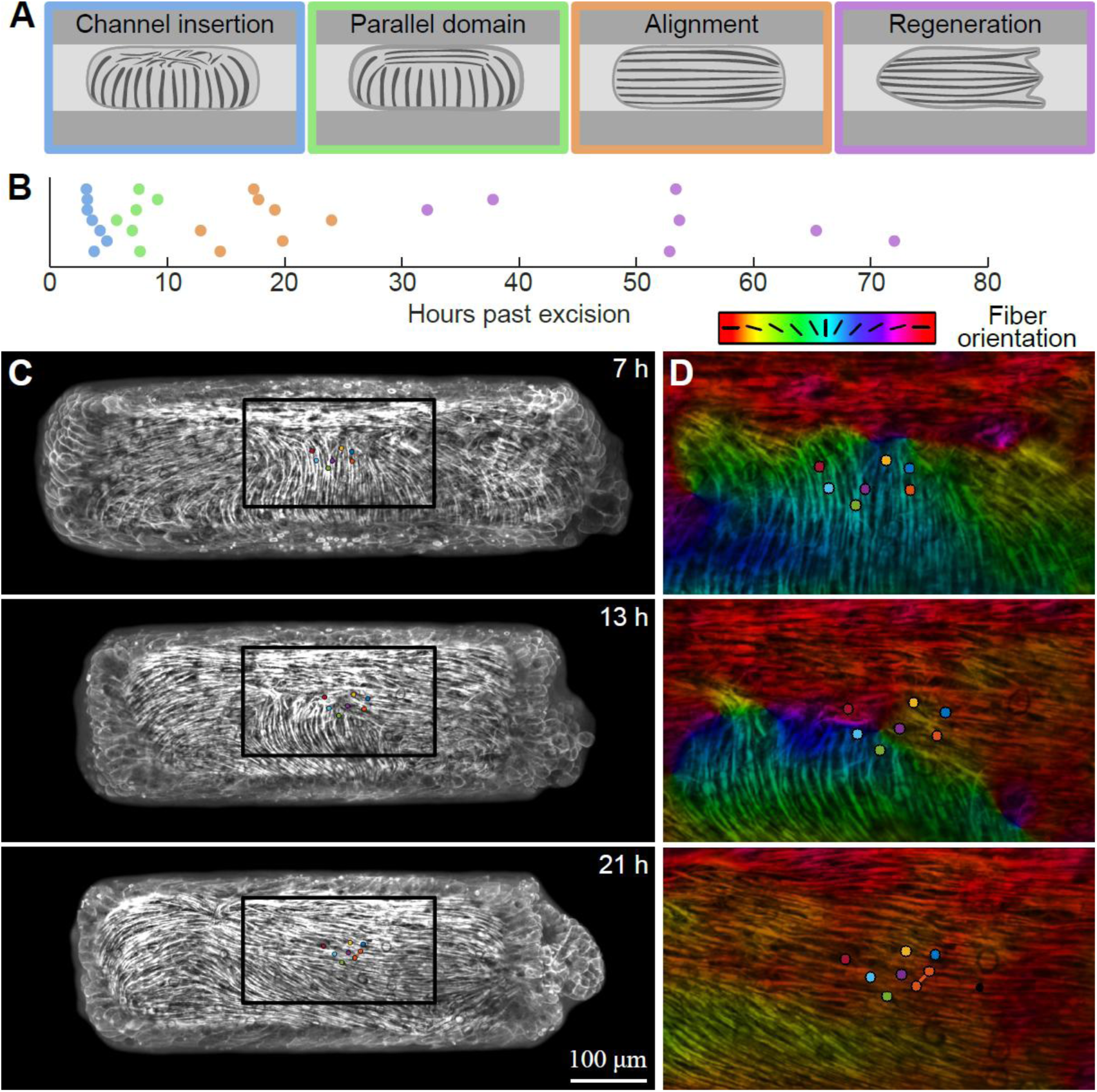
Reorientation dynamics of the actomyosin fibers. Schematics (A) and time-line (B) of the different steps in the reorientation of the actomyosin fibers in a confined tissue: (i) channel entry, (ii) formation of parallel fiber in the closure regions, with sharp domain boundaries with the inherited parallel domain (iii) establishment of a fiber array aligned with the channel axis, following the resolution of the extended domain boundaries, and (iv) regeneration (defined by the appearance of tentacles) as determined for 7 different samples confined in a perpendicular configuration (dots). (C) Projected images of the basal surface of the ectoderm showing consecutive stages of fiber organization in a regenerating Hydra spheroid expressing Lifeact-GFP confined in a perpendicular configuration (Movie 6). The images show the domain boundary between the parallel fibers formed in the closure region and the inherited fibers (top), expansion of the parallel domain following domain-boundary propagation (middle), and a later time point with fibers aligned primarily along the channel axis. (D) Zoomed views of the boxed region in C, color-coded according to the local fiber orientation (Methods). Due to the cyclic nature of the orientation field, we map the fiber orientation angle θ, relative to the channel axis in a clockwise manner (θ ∈ [0, π]) using a cyclic color map. The tracked cells from the apical surface are depicted. One cell divided, and the daughter cells are connected by a line.

**Figure 4.**
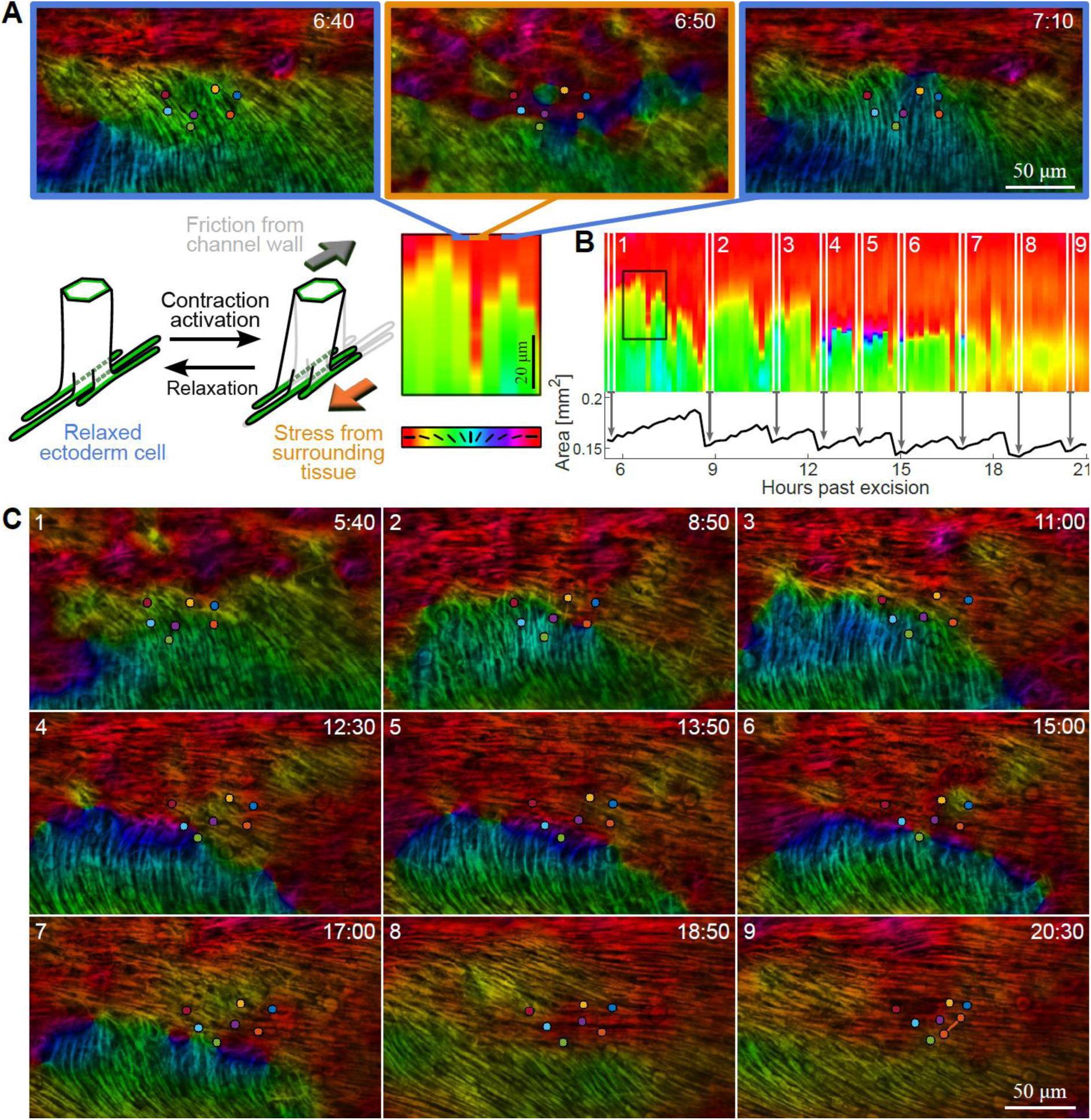
Propagation of the domain boundary. (A) A transient “jump” in the domain boundary position. Top: Zoomed images showing a transient “jump” in the domain boundary position, reflecting relative movement between the basal surface and the apical surface. The images show the fiber organization color coded according to their orientation as in Fig. 3D, before, during and after the transient “jump” of the domain boundary. Bottom left: scheme illustrating the relative movement between the apical and basal surfaces in an ectoderm cell upon transient contraction. Transient stress at the basal surface generated by widespread contraction of the ectodermal fibers displaces the basal side of the cell, whereas the apical surface remains nearly stationary due to friction with the channel wall. Bottom right: a zoomed view of the fiber orientation kymograph from the boxed region in B highlighting the transient “jump” in the domain boundary position. (B) Mean fiber orientation kymograph showing the reorientation dynamics. The kymograph is calculated from the boxed region in Fig. 3C, depicting the mean fiber orientation, color-coded as in Fig. 3D, as a function of time (x-axis) and position perpendicular to the channel axis (y-axis), after averaging over the horizontal axis within the boxed region (Methods). The trace of the projected tissue area over time is also shown (bottom). The white lines depict the post-rupture time points shown in C. (C) Images of the actomyosin fibers at the basal surface immediately after tissue rupture events in the zoomed region shown in Fig. 3D. At these post-rupture time points, following pressure release, the tracked cell positions at the apical surface serve as an approximate reference for the basal surface, showing that the domain boundary propagates in a directional manner relative to the tissue. The dots in A and C represent tracked cell positions in the apical ectoderm surface. One cell divided in C, and its daughter cells are connected by a line.

To characterize the movement of the domain boundary, we generate kymographs of the average nematic fiber orientation within a region of interest as a function of time (Fig. 4B; Methods). Across multiple samples and different viewing orientations, we obtain a consistent picture: the four extended domain boundaries, on both sides of the two original closure regions, propagate to expand the parallel domains and shrink the perpendicular domain. Propagation continues until, after roughly 10 to 20 hours of confinement, the two boundaries flanking each perpendicular domain meet, yielding the merger of the two parallel domains and the disappearance of the elongated domain boundary (Figs. 3, 4B,C). Eventually, the regenerated animal’s nematic organization exhibits only point defects, with no extended domain boundaries. Regenerated animals with normal morphologies typically have two defect regions, each with a net +1 charge, at either end of the elongated spheroid that will become the head and the foot of the regenerated animal, respectively. Samples which regenerate into abnormal morphologies have “excess” positive defects as well as negative - 1/2 defects (Fig. S2)^22^.

Movement of the domain boundary is largely directional: after its initial formation, the parallel domain expands at the expense of the perpendicular domain until the boundary disappears (Fig. 4). This propagation is intermittently interrupted by pronounced “jumps”, in which the boundary abruptly shifts and then relaxes toward its previous location (Fig. 4A). We attribute these jumps to transient fiber contraction; contraction of the perpendicular fiber domain, pulls neighboring tissue at the basal surface, displacing the domain boundary. Subsequently, upon relaxation, the boundary returns toward its prior position. This rapid contraction generates relative motion between the basal surface, where contractile forces are produced, and the apical surface, which remains nearly stationary, presumably due to friction with the channel wall (Fig. 4A). The transient jumps observed in the kymograph thus reflect tissue deformations, rather than motion of the domain boundary relative to the tissue. Over longer time scales, the domain boundary does move directionally relative to the tissue frame of reference. This propagation involves reorganization of the nematic fibers within the tissue, with tissue regions initially containing transverse fibers developing parallel fiber arrays aligned with the channel axis (Fig. 4C).

Global contraction events often lead to tissue ruptures, allowing luminal fluid efflux and the release of hydrostatic pressure (Fig. 4B)^12^. Interestingly, domain propagation appears more pronounced following rupture events, when tissue tension is abruptly reduced, suggesting that lower tension following ruptures may create more permissive mechanical conditions for fiber rearrangements (Fig. 4B,C).

To characterize the dynamics of the reorientation process with cellular resolution, we turn again to partially labeled chimeric samples (Fig. 5, Movie 7; Methods). By observing boundary propagation at the graft seam, we can visualize the reorientation process within individual cells. We segment the graft seam on the apical surface and track labeled cells over time (Fig. 5C, left). By overlaying the segmented apical seam on the basal surface, we can identify fiber protrusions at the basal surface that originate from adjacent labeled cells (Fig. 5C, right). We find that at the domain boundary, perpendicular fibers that protrude under neighboring cells above and below a given cell disappear. Subsequently after an intermediate period with no observable basal protrusions, parallel fibers reappear, protruding below different neighboring cells on either side of the cell (Figs. 5A,C, S8). Between the dissolution of the perpendicular protrusions to the formation of the parallel protrusion, the cells bordering the seam do not have observable parallel fiber arrays in the basal layer. These observations shed light on the mechanism of fiber reorientation, indicating that the domain boundary propagates by dissolution of the perpendicular fibers at the boundary and the reformation of fibers in a parallel orientation.

**Figure 5.**
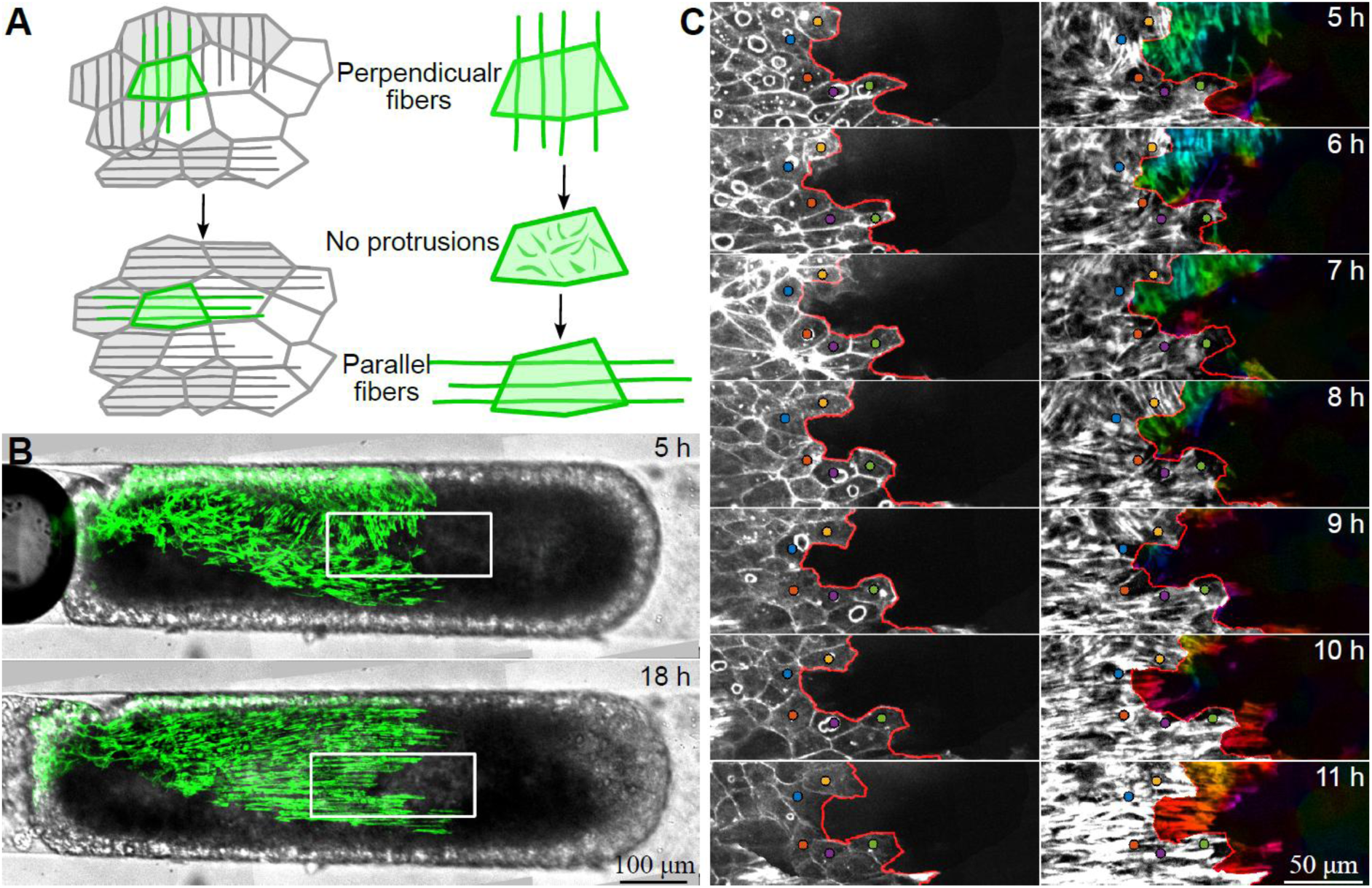
Fiber reorientation within individual cells. (A) Schematic illustration of fiber reorganization around the graft seam in a partially-labeled chimeric sample. Left: Initial and final fiber organization around the domain boundary. Cells that have perpendicular labeled protruding fibers initially (top) undergo reorientation and form parallel fibers (bottom). Right: Schematic illustration of the reorientation process within a single cell. Initially, the basal fibers are oriented vertically (top), subsequently the fibers dissolve and the cell becomes devoid of protruding fibers (middle) and finally fibers reform in a horizontal orientation (bottom). (B) Brightfield and fluorescence overlaid images of the initial (5 hours after excision, 1 hour after channel insertion) and reoriented (18 hours after excision) fiber organization in a confined partially-labeled chimeric sample. (C) Zoomed images of the boxed region in B, which is centered around the tracked cells, at 1 hour time intervals. Left: images of the apical surface of the ectoderm. Colored dots depict the position of manually tracked individual cells and the red line depicts the graft seam in the apical surface. Right: images of the basal surface of the ectoderm highlighting the fiber reorganization. The tracked cells and graft seam in the apical surface are indicated. The protruding regions, defined as regions beyond the red line, are color coded according to the local fiber orientation, as in Fig. 3D, highlighting the reorientation process.

## Discussion

In this work, we show that regenerating *Hydra* under frustrating confinement can reorient their body axis relative to the tissue frame of reference. Using high-resolution live imaging, we follow the nematic organization of supracellular actomyosin fibers and show that fibers form parallel to the channel axis within the initially disordered closure regions, creating abrupt domain boundaries with the inherited perpendicular fiber domains. These boundaries then propagate, so that both the primary fiber orientation and the regenerated body axis ultimately become aligned with the channel axis.

During reorientation, we observe the dissolution of the inherited, perpendicular basal fibers in cells adjacent to the domain boundary, followed by the formation of a thin, broad lamellipodium-like protrusion that subsequently develops into multiple parallel fibers (Fig. S8). Thus, the basic unit controlling fiber dynamics is individual cells, rather than individual fibers, since each cell contains several fibers. The interactions that promote dissolution of inherited fibers at the domain boundary and direct the orientation of newly formed fibers are still unknown.

Our observations show that the nematic fibers in confined *Hydra* tissues reorganize relative to the material frame, rather than through large-scale tissue flow. This behavior is consistent with previous observations from our lab showing that point defects in regenerating fragments are motile relative to the tissue^9^. The solid-like, elastic nature of the tissue is evident from its behavior under confinement, where the tissue sustains large anisotropic strains that persist over long time scales, without giving rise to large-scale tissue flow (Figs. S6,S7). Together, these observations support a framework in which *Hydra* tissues behave as active nematic elastic solids^12,23,28^. This contrasts with liquid nematics, where defect motion and director reorientation are strongly coupled to material flow^29,30^.

Having established that *Hydra* tissues behave as active nematic solids on the relevant timescale for axis reorientation, we next turn to the interactions that organize the nematic fiber-orientation field. Fiber-alignment interactions, which give rise to a nematic Frank energy, and are typically assumed to be the main mechanism underlying nematic order^12,29–32^, do not explain the observed nematic organization in confined tissues (see Supplementary Information). This is evident from the formation of parallel fibers in the closure regions, generating narrow domain boundaries with the adjacent inherited perpendicular fibers. If fiber-alignment interactions were dominant, fibers forming in the closure regions would be expected to connect smoothly with the inherited perpendicular fibers, as observed in numerical simulations of confined *Hydra* tissues^22^. Our observations further show that the domain boundaries remain abrupt as they propagate through the tissue over hours. This indicates that the domain boundaries are not simply quenched at their initial positions, but rather are dynamically sustained as the system develops. Together, these observations imply that nematic order in *Hydra* cannot be explained by local fiber-alignment interactions alone and must also involve coupling to additional external or internal fields (Supplementary Information).

Under confinement, the tissue is stretched along the channel axis. New fibers, whether formed early in the wound-closure regions or later during domain-boundary propagation, align with the channel axis. The coincidence between the orientation of the tissue strain and the preferred nematic fiber orientation strongly suggests that fiber orientation is biased by the primary strain direction in the tissue. Similarly, planar confinement between two surfaces was shown to bias the orientation of the regenerated body axis and to promote the formation of biaxial *Hydra*^23^. These observations are also consistent with recent work on regenerating *Hydra* cell aggregates, showing that anisotropic stretch biases newly formed fibers along the direction of stretch^14^. Importantly, the magnitude of externally induced strain in these examples is comparable to naturally occurring tissue deformations generated by active actomyosin contractions^12^. As such, strain coupling could provide mechanical cues that direct fiber orientation in regenerating *Hydra*, complementing local alignment interactions between neighboring fibers (Supplementary Information)^28^. In this picture, activation of contraction along the nematic actomyosin fibers generates tissue strain, which in turn feeds back on fiber orientation and remodeling.

Although we focus here on strain coupling to the nematic, the orientation of the fiber network may also be influenced by other internal and external cues, such as tissue curvature, or coupling to biochemical concentration fields or electrical fields^33,34^. Surface curvature was shown to be coupled to orientation in nematic elastomers^35^. Indeed, confinement in narrow channels introduces clear anisotropy in tissue curvature. However, previous work on *Hydra* aggregates, considered samples with a similar elongated geometry but without the associated strain pattern, and did not observe a bias in fiber orientation^14^, suggesting that coupling between nematic orientation and surface curvature is not a dominant factor. Another possibility is interaction between the nematic field and biochemical fields. Previous theoretical work by us and others showed that alignment to gradients of a scalar morphogen concentration field can help stabilize +1 defects and recapitulate many aspects of *Hydra* regeneration^12,32^. Given the initial radial symmetry of the excised ring and the sealed tissue spheroid before confinement, it is unlikely that a strong pre-existing signaling gradient is initially aligned with the channel axis. Nevertheless, signaling gradients could arise in response to confinement and subsequently influence nematic remodeling. Altogether, our results support a picture of an active nematic elastomer, in which the nematic fiber organization emerges from coupling between local fiber remodeling, tissue strain, and potentially other signaling processes. How these different cues are integrated across spatial and temporal scales remains a central open question.

The primary axis of fiber orientation and the body axis in *Hydra* are always aligned. Under unperturbed conditions, excised tissues regenerate along their inherited primary fiber orientation, aligned with the body axis of their parent animal^11,16^. In contrast, here we show that regenerating tissues can reorient their body axis in response to mechanical constraints, relative to both the inherited body axis and the initial primary fiber orientation. This suggests that mechanical features such as the nematic fiber orientation and the strain pattern are coupled to biochemical signals, which together give rise to body axis formation through synergistic interactions^12,36^. The proposed ability of cells to locally align fibers with the direction of strain could close a mechanical feedback loop: tissue-scale strain biases local fiber orientation, while the resulting fiber organization shapes force generation and mechanical integration across the tissue, ultimately giving rise to long-range fiber organization and body axis formation. Such feedback between tissue mechanics, cytoskeletal organization, and axis formation is likely to be broadly relevant to morphogenesis in other organisms. A conceptually related, but mechanistically distinct, case is found in plants, where cortical tension guides the orientation of cortical microtubules, which in turn bias anisotropic cell-wall deposition and growth, providing another example of mechanically guided cytoskeletal reorganization shaping morphogenesis^37^. The ability to induce body-axis reorientation externally, while following the process by high-resolution live imaging, provides a powerful platform for testing this feedback in animals and examining how the interplay between mechanical, electrical, and biochemical fields orchestrates body-axis patterning.

## Acknowledgments

This work was supported by a grant from the Israel Science Foundation to K.K. (grant No. 3565/24). This work was supported by the Institut Ruđer Bošković Program “Natječaji za jačanje potencijala mladih znanstvenika”, thematic component “Podrška povratnicima” (MZ1-26), decision number 01-1106/2-2026 to M.P., funded by the European Union – NextGenerationEU. We thank all the members of our lab for help, discussions and comments on the manuscript. We thank Yonit Maroudas-Sacks, Erez Braun, Anais Ballias, S Suganthan, Fridtjof Brauns, Jana Fuhrmann, Suat Ozbek and Ram Adar for discussions and comments on the manuscript.

## Methods

### Hydra strains and culturing

Experiments were performed with two transgenic strains of *Hydra* vulgaris (AEP): a strain expressing Lifeact-GFP in the ectoderm^38^, generously provided by Bert Hobmayer, University of Innsbruck; and a strain expressing EGFP-tagged Hyβ-catenin in the ectoderm developed by the Holstein lab^19^, University of Heidelberg, and generously provided by Charisios Tsiairis, Friedrich Miescher Institute for Biomedical Research, Basel, Switzerland. Animals are cultivated in Hydra medium (HM; 1mM NaHCO3, 1mM CaCl2, 0.1mM MgCl2, 0.1mM KCl, and 1mM Tris-HCl, pH7.7) at 18°*C*. The animals are fed with live Artemia nauplii three to five times a week and subsequently washed ∼6-10 h after feeding.

### Grafting to generate chimeric-labeled Hydra animals

Grafting of different transgenic strains is used for generating chimeric-labeled *Hydra* for stable tissue tracking (Fig. 1C) and for visualizing the dynamics of individual ectodermal actin fibers at the graft seam (Fig. 5). Grafting is done as previously reported by axial grafting the lower half of a bisected animal from one transgenic line to the upper half of a bisected animal from a different transgenic line^16,39^. Grafting is done using a strain expressing Lifeact-GFP in the ectoderm that allows visualization of the actin organization and a strain expressing EGFP-tagged Hyβ-catenin that appears unlabeled in the imaging conditions used (due to the inherently low expression levels in this strain). Grafted animals are fed and typically used 2-5 days after grafting. Partially labeled buds emerging from grafted animals sometimes have a graft seam roughly aligned with their body axis. Such a bud was used to generate the sample shown in Fig. 5.

### Sample preparation

Tissue excision is done ∼24 hours after feeding. Tissue rings, about 1/5 to 1/4 of the original body length. are excised from the mid-body section of an adult *Hydra*. This tube length is large enough to avoid buckling when folding, yet short enough to form an oblate spheroid that will align with its symmetry axis perpendicular to the flow during confinement. Excision of rings from axially-grafted animals is done in a similar manner, making sure that the transverse cuts are done around the graft seam to form a differentially labeled tissue ring.

### Confinement in cylindrical hydrogel channels

We embed regenerating *Hydra* tissue in cylindrical channels made of agarose gel as described previously^22^. The channel walls are made of 2% gel. The channels are prepared using a pulled glass capillary with a tapered neck and long, narrow head of approximately 180-300 μm in width. The glass capillaries are placed in a custom-made mold with a narrow slit designed to hold the glass capillary in place, so that the tapered, narrow end of the capillary is suspended in a hollow cavity of around 0.5 cm in diameter. The cavity is then filled with liquefied 2% agarose gel, which sets around the glass capillary. Once the gel has set, the capillary is removed by pulling backwards leaving a hollow cavity in the gel in the shape of the capillary. The same capillary is then truncated to remove the narrow end, and re-inserted into the gel, so the tapering end is aligned with the narrowest part of the channel in the gel and creates a funnel leading into the channel. A small hole is made from above using a narrower capillary at the far end of the channel, to prevent pressure build-up in the channel during sample insertion. The *Hydra* tissue spheroids are then inserted into the channels by flowing them in a media of liquefied 0.5% low melting agarose gel through the glass capillary and into the channel using a syringe. The inserted tissue spheroid becomes lodged in the narrow end of the channel, and the soft gel that sets around it inside the channel and prevents it from flowing back into the wide section of the channel. The full agarose block containing the channel is removed from the mold, and excess gel is removed by cutting around the sample in the channel, to allow more samples to be placed together in the imaging dish. Multiple agarose gel blocks, each containing a confined sample, are mounted in a glass-bottomed dish for imaging. The mounting is secured with liquefied 0.5% gel that stiffens and serves as a glue.

During the insertion process, the sample is exposed to large shear forces. These shear forces induce tissue rupture. This is apparent by eye from the decrease in the spheroids’ volume before and after insertion, as well as from following the saw tooth pattern of the projected area of the regenerating tissue after insertion into the channel, which typically initiates at a low value, characteristic of the relaxed, post-rupture tissue state (See the extrapolated area in Fig. S1). The ruptures induced during insertion into the channel appear similar to the spontaneous tissue ruptures that normally occur as part of the osmotic inflation-deflation cycles of regenerating *Hydra* tissues, associated with a rapid decrease in lumen volume and relaxation of tissue tension. In most cases these ruptures heal within minutes^12^. In some cases, the insertion process can lead to large tears in the tissue, in which the damage to the integrity of the tissue is still apparent once we start imaging 1.5-3 hours after channel insertion, or we observe excessive sample shrinking due to extrusion of dead cells during the first hours of imaging. These samples were excluded from the experiments, and in all subsequent analyses we consider only tissues that appear intact.

We further filter the confined samples as follows. The initial orientation of the principal fiber orientation relative to the channel is classified as aligned with the channel, or transverse to the channel. This was judged based on the primary orientation of the inherited, ordered supracellular fibers. Importantly, this was judged “blind” per sample, without knowledge of the final outcome. Since the samples undergo osmotic inflations while maintaining a constant width defined by the channel diameter, each sample’s aspect ratio varies over time. To obtain a well-defined measure, the aspect ratio is determined from the frame immediately following the first tissue rupture, identified as the first large, rapid drop in tissue area. For all further analysis, we consider only samples lodged in the transverse configuration and elongated so that their long axis is at least twice their diameter.

To observe the outcome morphology of samples, the tissues are released from the channels and imaged without external confinement. The gel block is placed in HM, and a transverse cut is made on the wider side of the channel, perpendicular to the channel’s axis. This cut is positioned as close as possible to the sample, shortening the channel without causing damage. Plastic forceps are then used to gently “squeeze” the sample out of the channel by pressing on the gel block from the side of the sample opposite to the newly created exit (Fig. S7).

### Tissue labeling using photoactivation of caged dyes

To label specific tissue regions we use laser-induced activation of a caged dye (Abberior CAGE 552 NHS ester) that is electroporated uniformly into mature *Hydra* and subsequently activated in the desired region as described previously^9^. Electroporation of the probe into live *Hydra* is performed using a homemade electroporation setup. The electroporation chamber consists of a small Teflon well, with 2 perpendicular Platinum electrodes, spaced 2.5 mm apart, on both sides of the well. A single *Hydra* is placed in the chamber in 10 µl of HM supplemented with 2 − 12 mM of the caged dye. A 75 Volts electric pulse is applied for 35 ms. The animal is then washed in HM and allowed to recover for several hours to 1 day prior to tissue excision. Activation is done on samples after insertion into the channel. The specific region of interest is activated using a 100 mW 405 nm laser in a spinning disk confocal system equipped with a Vector fixed point laser photobleaching system (3i, Intelligent Imaging Innovations) using a 10× air objective (NA=0.3). Subsequent imaging of the Lifeact-GFP signal and the uncaged cytosolic label is done by spinning-disk confocal microscopy as described below.

### Microscopy

Spinning disk confocal movies of regenerating *Hydra* are acquired in one of two systems (Intelligent Imaging Innovations) running Slidebook software. The first system is an inverted spinning-disk confocal microscope. Lifeact-GFP is excited using a 50 mW 488 nm laser and the activated Abberior CAGE 552 is excited using a 50 mW 561 nm laser. Time-lapse movies are acquired using an EM-CCD (QuantEM; Photometrix) using a 10× air objective (NA = 0.5). The second system is an upright spinning-disk confocal microscope. The Lifeact-GFP is excited using a 200 mW 488 nm laser. Time-lapse movies are acquired with a sCMOS camera (Andor Zyla 4.1) using a 10× air objective (NA=0.3) or a 20× dipping objective (NA=0.5) from above. Since imaging from above required an open dish, perfusion of media is employed to compensate for evaporation.

In both systems, z-stacks are taken with a z-resolution of 3 μm, and a time interval of 2-10 minutes over 2-3 days, until regeneration is complete. For each time point, brightfield images are acquired at the sample mid-plane, together with spinning-disk confocal z-stacks. While the channel width is small enough to fit within the field of view, most samples are too long to fit in a single field of view. To image the entire sample, several positions are defined for each sample, with partial overlap between adjacent fields of view, to facilitate image stitching as described below.

For most time-lapse imaging experiments, individual samples are removed from the channels 3 days after excision, and imaged to obtain clear images of their final morphology. For this, animals are relaxed in 2% urethane in HM for 1 minute and then sandwiched between two glass coverslips with a 200 μm spacer between them. The final images are taken from both sides by flipping the sample.

### Image processing and analysis

#### Image stitching

Multiple (typically 2-3) images of adjacent regions of the same sample, overlapping by approximately 50-100 μm, are stitched to show the entire sample using the Pairwise Stitching plugin in ImageJ (plugin internal version 1.2) ^40^. Stitching is performed on 2D maximum intensity projections for the first time point of each acquisition segment and applied to all subsequent frames. Alignment is computed using the built-in phase correlation algorithm, allowing translational shifts only. Fused images are generated using maximum intensity fusion.

#### Creating masks of tissue region

To define the tissue region for further analysis and determine properties such as the projected area, tissue length and width, masks are generated based on the max intensity projections of the Lifeact-GFP signal, using automatic thresholding in ImageJ. “Li” method is used in most cases, but depending on the quality of the image other automatic thresholding are used, and subsequently smoothed.

#### Surface detection and layer separation

The curved apical and basal surfaces of the ectoderm, containing the apical cell-cell junctions and the basal actomyosin fibers respectively, are computationally identified in the 3D spinning disk z stacks of regenerating spheroids expressing Lifeact-GFP in the ectoderm using the “Minimum Cost Z surface Projection” plugin in ImageJ as previously described ^12^. The cost images are generated by preprocessing the original z-stacks using a custom code written in MATLAB. First, the signal from the ectoderm layer is manipulated to make it more homogeneous within the layer without increasing its thickness, by applying the built-in MATLAB anisotropic diffusion filter. Then, we apply a gradient filter to highlight the apical and basal surfaces as the top and bottom boundaries of the ectoderm layer. The apical and basal surfaces are determined using the minCost algorithm (Parameters used: rescale xy 0.25; rescale z 1; min distance, 15 μm; max distance, 45 μm; max delta z, 1; two surfaces). The surfaces identified by the minCost algorithm are smoothed by applying an isotropic 3D Gaussian filter of 1-3 pixels in width (after rescaling to an isotropic grid matching the resolution in the z-direction) and selecting the iso-surface with value 0.5.

#### Image projection

2D projected images of the ectodermal actin fibers in the basal surface, and the cellular cortices in the apical surface, are generated by extracting the relevant fluorescence signal from the 3D spinning disk confocal image stacks based on the surface determined above. To determine the projected value for each x-y position, a Gaussian weight function in the z-direction with a sigma of 3 µm, centered at the z-value determined from the smoothed surface with a small (2–3 pixels) fixed offset found to optimally define the desired surface is employed. To produce 2D images of the cells and fibers, an offset range that best showed the signal for the basal/apical surface is manually chosen and a maximum intensity projection of these surfaces is created. The resulting 2D projected images are further subject to contrast limited adaptive histogram equalization (CLAHE) with MATLAB’s “adapthisteq” function with Rayleigh distribution and a tile size of 26 µm and the binary mask is applied again to the adjusted images. For samples that are imaged in multiple tiles, surface detection and layer separation is done separately on each stack. The resulting projection images are stitched by recomputing rigid translational registration using the parameters determined for the corresponding maximum intensity projections as described above.

In images of chimeric animals, detection of the apical surface around the graft seam is often poor. Better projections of the graft seam region in the apical surface, in the central part of the image are generated by creating a maximum z-projection of a subsection from original stack, containing the apical signal but not the basal. This method is limited as it does not produce proper projections over the entire sample but is useful for creating projections in the central part of the channel as shown in Fig. 5.

#### Actin fiber orientation analysis

The local orientation of the ectodermal supracellular actin fibers can be described by a nematic director field, oriented along the mean orientation determined in a small region surrounding every point. The director field is extracted in an automated way from the 2D projected images of the fibers as previously described ^9^. Images showing fibers color coded according to their orientation are created using the “hsv2rgb” function in MATLAB, using the normalized grayscale image as the Value matrix, the orientation field as the Hue, and the Saturation is set to 1.

For orientation kymographs, we manually chose a rectangular area on the stitched movie that contains both parallel and perpendicular domains at the beginning of the movie. The vertical length of the rectangle determines the kymograph’s vertical scale, and the mean orientation for each time point and each vertical position is calculated by averaging the orientation field across the horizontal axis within the rectangle. Concretely, the average orientation angle *θ_a_* at each vertical position is determined as

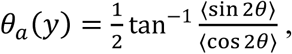

where *θ* is the local nematic orientation angle, and the averages ⟨sin 2*θ*⟩ and ⟨cos 2*θ*⟩ are evaluated across each horizontal stripe.

#### Tissue tracking

To identify the tissue frame of reference several methods are used, depending on the type of sample and the imaging setup. In samples labeled using a photoactivated dye, a long stripe of tissue is activated along the long axis of the confined tissue. The labeled tissue stripe is then followed over time by following bleach-corrected, maximum-projection images after applying a median filter with a ∼10 µm window size.

In samples imaged using a 20× dipping lens, the tissue frame of reference was tracked by manually tracking individual cells on the apical surface projection images. For the montage in Fig. 5, the region of interest was defined relative to the center-of-mass (COM) of the tracked cell positions. The COM trajectory was smoothed using a Savitzky-Golay filter and used to define a region of a fixed size around the COM, which was extracted from the projected apical and basal surface projections at each time point. Detection of the graft seam in the apical surface of grafted samples was done manually by segmenting the labeled region.

#### Statistical analysis

For analysis of the initial fiber orientation in the closure region following confinement in the perpendicular configuration within channel (for samples with an aspect ratio >2; see above), a pool of N=38 samples expressing LifeactGFP was considered. Out of these samples, 3/38 were facing the objective so that only inherited fibers were visible and the fiber orientation in the closure region could not be determined (Fig. S5B), but in all other 35 samples, a parallel domain formed in the initial closure region, with a visible domain boundary between the inherited perpendicular fibers and the newly-formed parallel fibers in the closure region. The time of parallel domain formation was determined manually by observation. In most of the samples (N=24/35) the parallel domain was clearly visible already at the onset of imaging (1-3 hours after insertion into channels), implying it formed earlier.

For analysis of reorientation of the inherited actin fiber array, only samples that had clear tracking of the tissue frame of reference and were stably confined in the channel up to >24 hours after excision are considered. Tracking of the tissue frame of reference was possible in samples that were either imaged in high magnification, allowing manual tracking of cells on the apical surface (N=18), or samples that were images in lower magnification but had a fluorescent photoactivated tissue marker (N=13). Overall, in N=31 samples we could observe the inherited fibers and reliably track the tissue frame of reference for at least 24 hours. Out of those, in 3/31 cases the tissue rotated within the channel within the first 24 hours of the experiment. In all remaining 28/31 confined samples the tissue was stably confined and did not rotate, and we observed reorientation of the fibers in the tissue’s frame of reference, from the initial transverse orientation of the inherited fibers to a parallel orientation.

Analysis of the timing of the different stages of the reorientation process in Fig. 3B, is manually determined by observing the basal surface projection of N=7 samples imaged in high magnification and stably confined in the channel for the first 24 hours.

The regeneration outcome of samples confined in a perpendicular configuration and released from the channels after 3 days is classified into categories based on the number of feet/heads and any other protrusions observed, and how they are arranged relative to each other. The presence of a head and foot is judged based on tissue morphology. The different morphological outcomes are classified as follows: normal morphology (single head and foot along a single axis), abnormal morphology (a single head and foot, with an additional ectopic tentacle, foot or foot-like protrusion on the body), multi-headed morphology (two heads), stuck (alive but with no observed tentacles or hypostome), or dead (disintegrated).

## Supplementary Information

### Supplementary Figures

**Figure S1.**
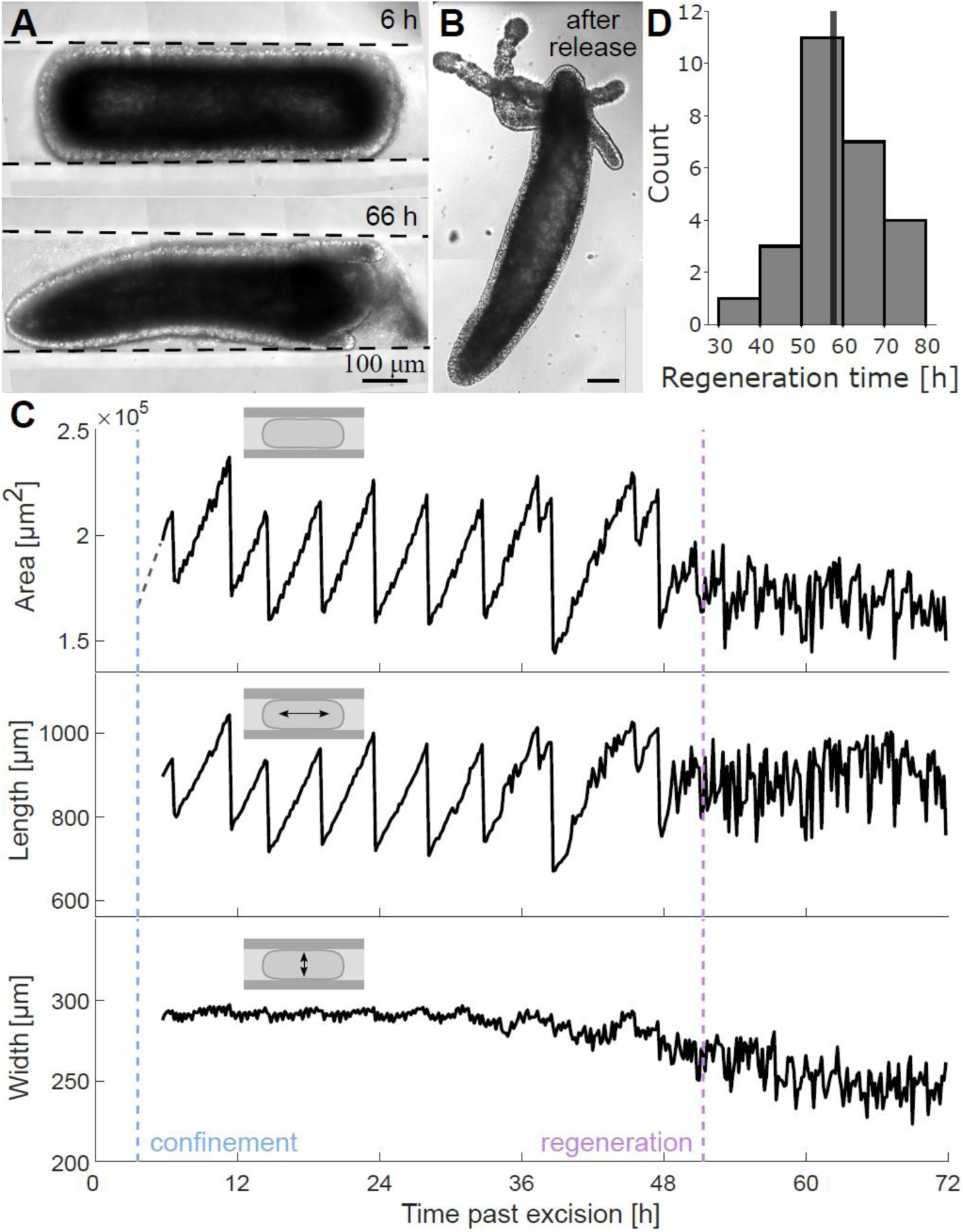
Regeneration in channels in the frustrated perpendicular configuration. (A) Brightfield images of a regenerating *Hydra* confined in a channel 6 hours past excision (∼2 hours after channel insertions) and 66 hours after excision. Tentacles are visible at 66 hours. (B) Brightfield image of the same regenerated *Hydra* after it was removed from the channel, showing the final morphology. (C) Traces of the projected area (top), length along the channel axis (middle) and width along the transverse axis (bottom) extracted from a time-lapse movie of the sample shown in A (Movie 1). The blue and purple dashed lines depict the time of channel insertion and regeneration (determined by tentacle appearance), respectively. Since the sample is constrained by the channel walls in the radial direction, the sample width is effectively set by the diameter of the channel and hardly changes over time. Only at later stages of the regeneration, when the animal starts to elongate along its regenerated body axis, the width of the sample becomes smaller than the channel diameter. In contrast, the length of the sample along the channel’s axis can vary freely and we observe similar saw-tooth pattern in the length trace over time as in the projected area. The dashed gray line in the projected area trace shows extrapolation of the first saw-tooth to the channel entry time. (D) Histogram of regeneration times for confined samples in the perpendicular configuration (N=26).

**Figure S2.**
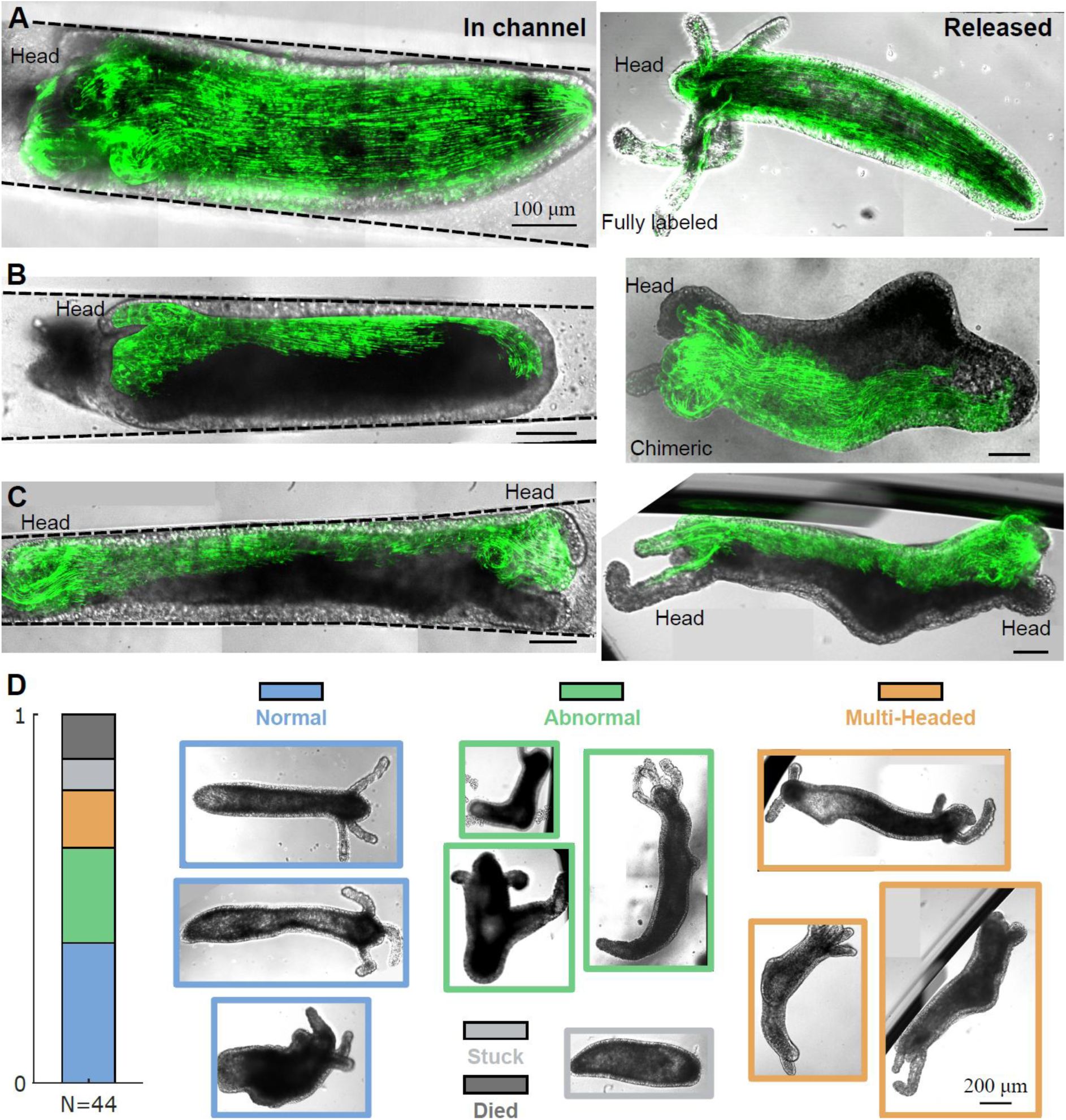
Regeneration outcomes of confined spheroids in the frustrated perpendicular configuration. (A-C) Brightfield and fluorescence overlaid images of regenerated *Hydra* after reorientation, before (left) and after (right) release from confinement. (A) Lifeact-GFP expressing animal with normal morphology. (B) Chimeric animal with abnormal morphology. (C) Chimeric animal with multi-headed morphology. (D) Left: Statistics of regeneration outcomes for tissue rings confined in a perpendicular orientation (with no appreciable tissue rotations for 24 hours). Right: Bright field images showing examples of regenerated *Hydra* with the different regeneration outcomes including normal morphology, abnormal morphologies with one head, multi-headed abnormal morphologies and stuck samples (see Methods)

**Figure S3.**
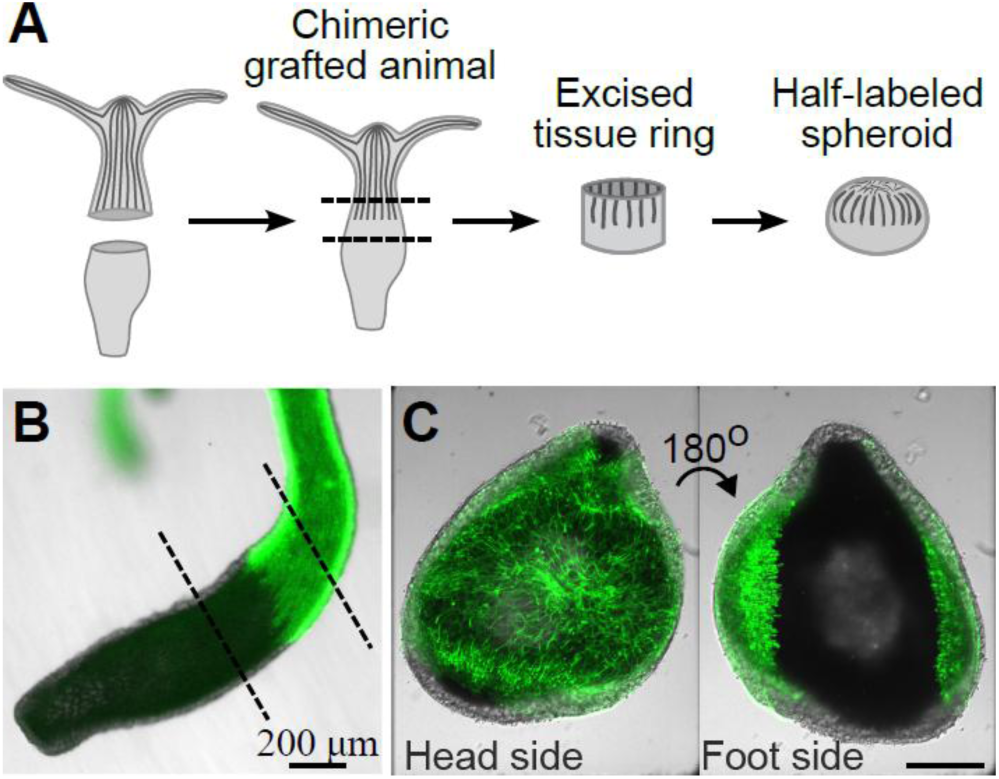
Axial grafting of chimeric *Hydra*. (A) Schematic illustration of tissue grafting and excision of a tissue ring from around the graft seam. (B) Brightfield and fluorescence overlaid image of part of a chimeric animal generated by grafting the head-half of an animal expressing Lifeact-GFP in the ectoderm with the foot-half of an animal that does not. (C) Brightfield and fluorescence overlaid images of opposite sides of a half-labeled spheroid, formed from a tissue ring excised from the middle of a chimeric animal, around the graft seam. The left image shows the closure region on the originally head-facing side of the ring which is labeled, and the right image shows the closure region on the opposite side which is unlabeled.

**Figure S4.**
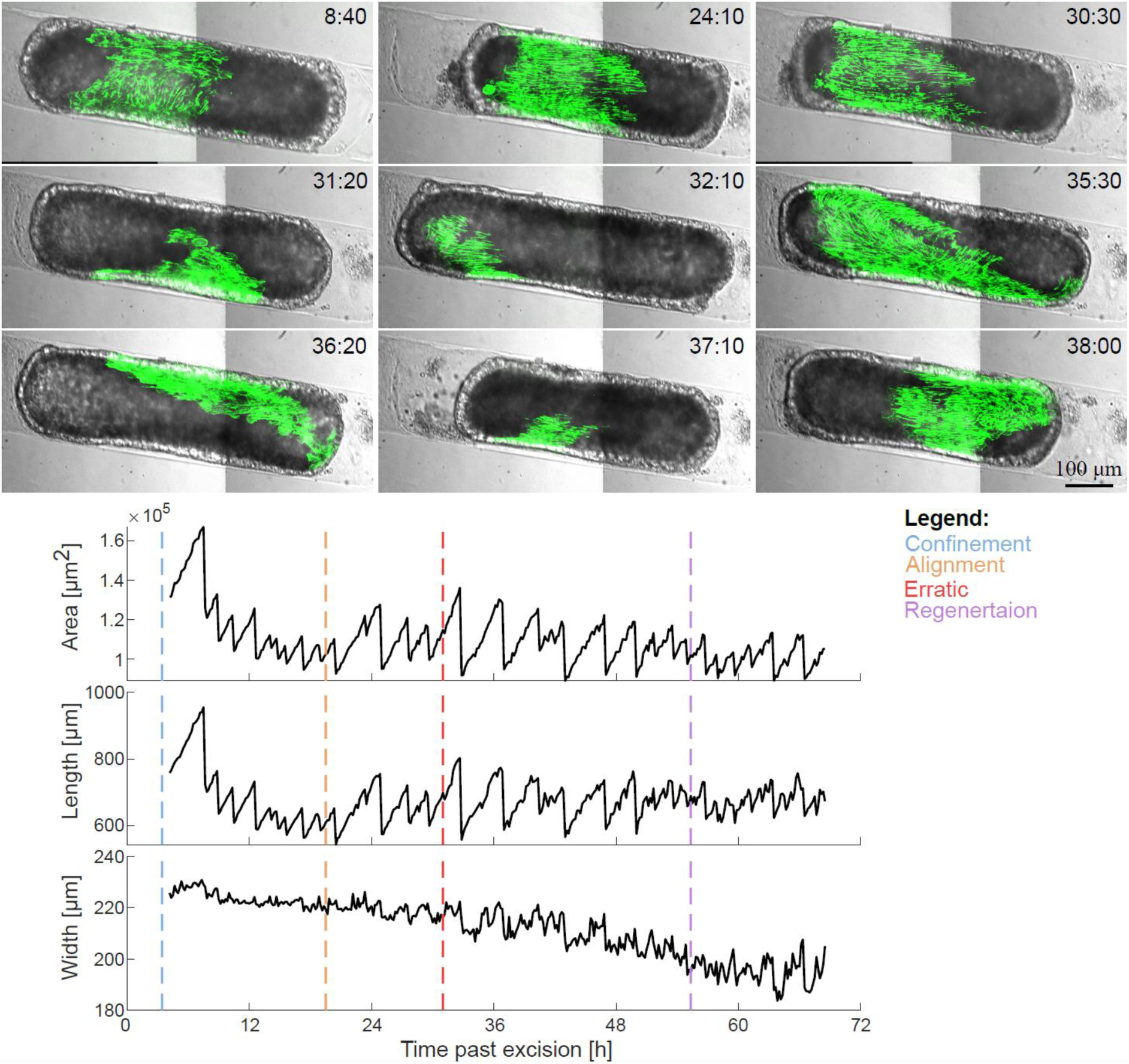
Transition from stable confinement to erratic movement within the channel. (A) Brightfield and fluorescence overlaid images of a partially-labeled chimeric sample confined in a channel (Movie 5). The tissue is stably confined for the first 30 hours past excision, as evident from the position of the graft seam. During this time, the inherited transverse fibers (apparent in the first image) reorient relative to the tissue (first row). Around 31 hours past excision, the sample undergoes an abrupt transition and starts rotating erratically in the channel. (B) Projected area, length and width of the regenerating *Hydra* shown in A over time. The dashed lines indicate different steps in the process: blue- channel entry; orange-alignment of fibers with the channel axis; red- transition to erratic rotations in channel; purple-regeneration (defined by tentacle appearance).

**Figure S5.**
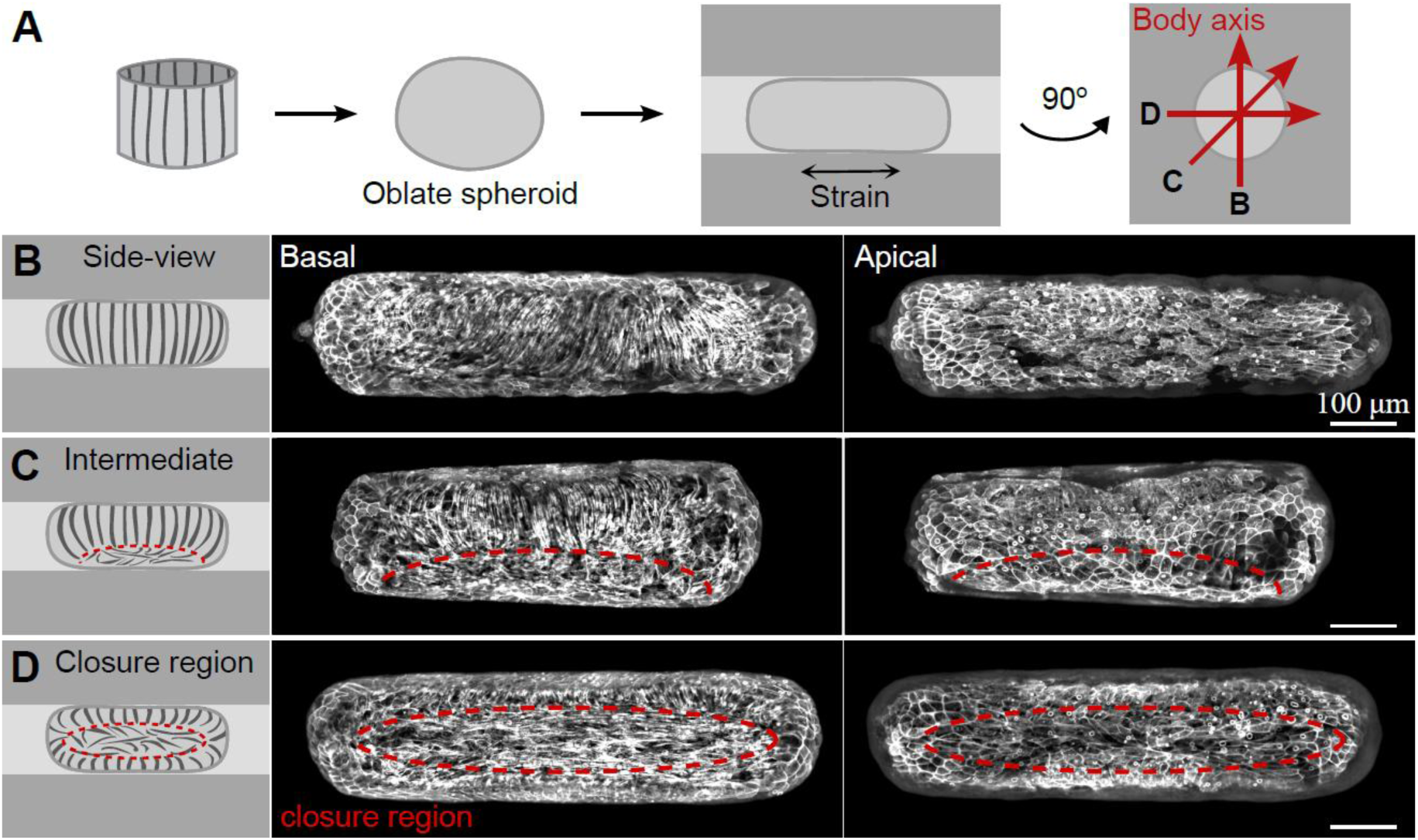
Examples of different viewing orientations in the frustrated perpendicular configuration. (A) Schematic illustration of the experimental setup: an excised tissue ring seals into an oblate spheroid that is inserted into a channel at different possible viewing orientations. (B-D) Scheme (left) and images of the basal and apical surfaces of the ectoderm for confined samples in the perpendicular configuration with (B) a complete side-view showing only inherited fibers transverse to the channel axis; (C) Intermediate view, where the tissue facing the objective consists of inherited transverse fibers (top), and one of the closure regions with disordered fibers (bottom); and (D) Head-on view of one of the closure regions, which is surrounded by inherited fibers which are visible around its edge.

**Figure S6.**
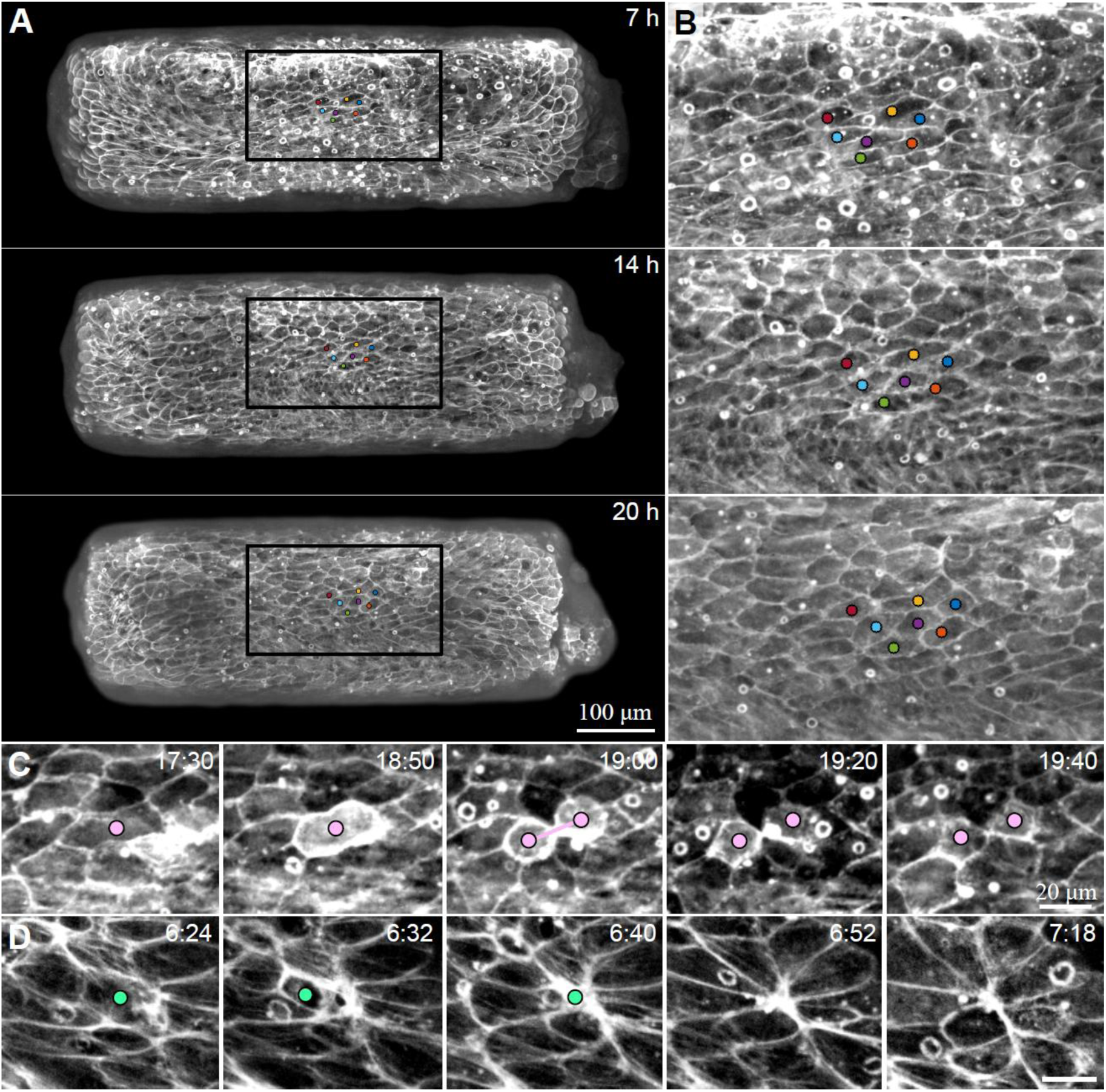
Cell shape anisotropy in confined spheroids. (A) Images of the apical surface from a movie of a confined *Hydra* tissue expressing Lifeact-GFP in the ectoderm at 7, 14 and 20 hours past excision (confinement at 3 hours past excision). Colored dots depict manually tracked individual cells. (B) Zoomed images of the boxed region in (A). Note that the cell shapes are still elongated along the channel axis 20 hours past excision. (C) Sequential images showing a cell division event. (D) Sequential images showing a cell extrusion event.

**Figure S7.**
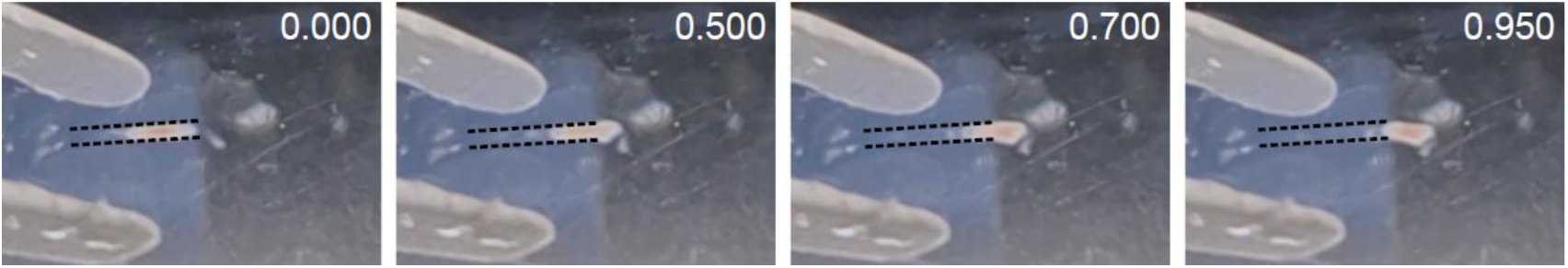
Fast recoil of the tissue upon release from confinement. Consecutive images of a sample released from the channel 3 days after confinement. Dashed lines denote the channel walls. The images show the tissue recoils and become wider as soon as it exits the channel, indicating that the confined tissue stores strain even 3 days after being placed in the channel. The time during the release process is indicated in fractions of a second.

**Figure S8.**
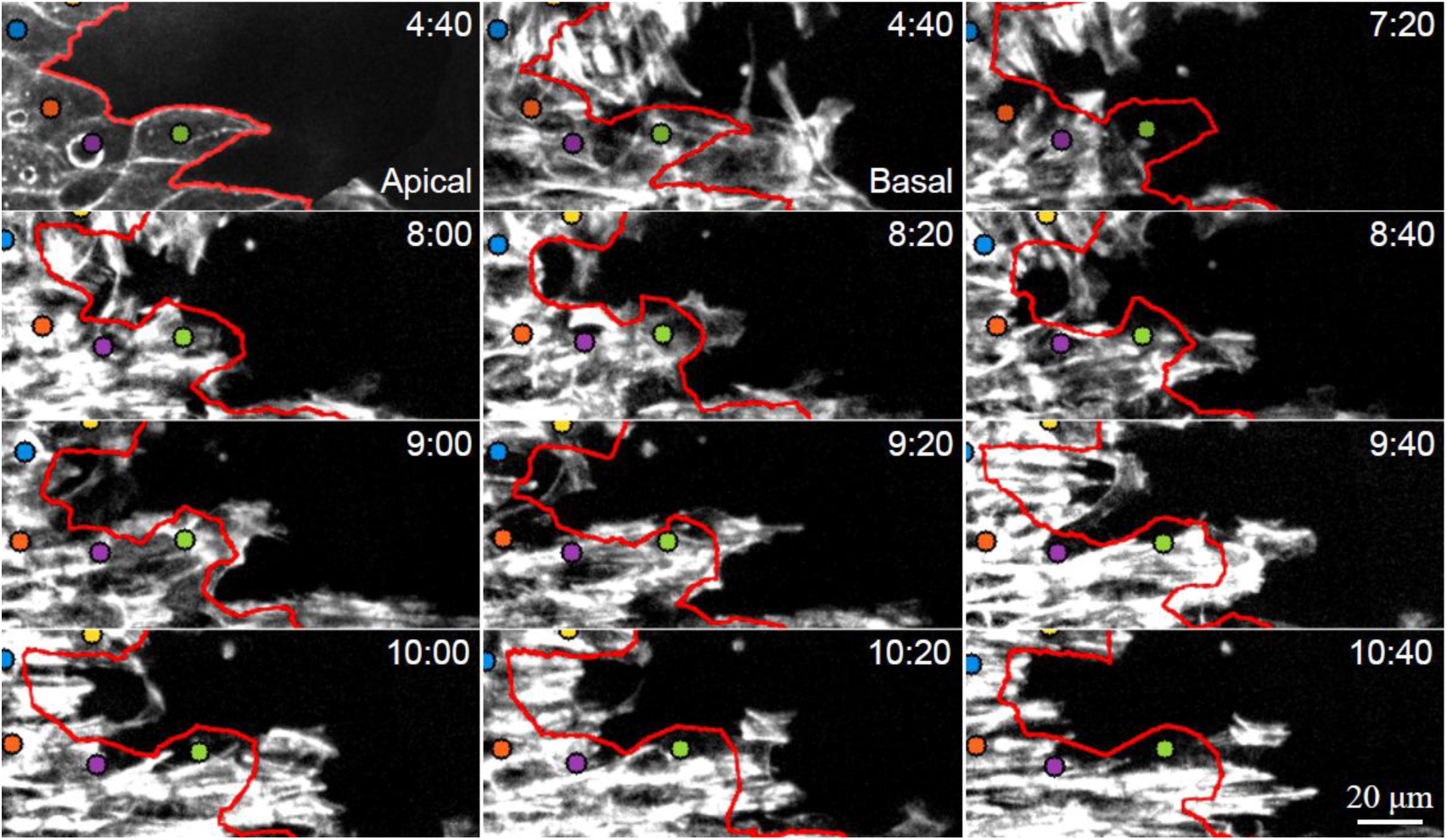
Formation of new fiber protrusions at the domain boundary. Images from a time-lapse movie of a confined, partially labeled chimeric regenerating *Hydra* (Fig. 5, Movie 7) showing the formation of fiber protrusions near the cell labeled with a green dot. The first image shows the apical surface, and all subsequent images show the basal surface at consecutive time-points. The images show the growth of a thin, broad lamellipodium-like protrusion preceding the formation of parallel protruding fibers.

### Supplementary Movies

**Movie 1: Bright-field movie of a regenerating *Hydra* spheroid confined in a perpendicular configuration.** Bright-field time-lapse movie of a regenerating *Hydra* spheroid under confinement (Fig. S1). The tissue was excised and allowed to seal into an oblate spheroid, before being inserted into a narrow cylindrical channel using flow, 3.5 hours after excision (Methods). The geometry and shape dynamics of the regenerating tissue are constrained by the channel, which limits the width of the confined spheroid as it deforms. The confined tissue spheroid undergoes cycles of osmotic inflations and rapid deflations, and eventually regenerates into a mature *Hydra*, which has a single body axis aligned with the channel. The elapsed time from excision is displayed (hh:mm), and the scale bar is 100 µm.

**Movie 2: Regenerating *Hydra* spheroid confined in a perpendicular configuration labeled with a photoactivated dye.** Time-lapse, spinning-disk confocal movie of a confined regenerating tissue spheroid expressing Lifeact-GFP in the ectoderm. The tissue was introduced into the channel in a perpendicular configuration 5.5 hours after excision and then labeled with a photoactivated tissue marker (Abberior CAGE 552; Fig. 1B). Projected images show the Lifeact-GFP signal (gray) overlaid with the photoactivated tissue marker (magenta). The elapsed time from excision is displayed (hh:mm), and the scale bar is 100 µm.

**Movie 3: Regenerating chimeric *Hydra* spheroid confined in a perpendicular configuration.** Time-lapse, spinning-disk confocal movie of a chimeric tissue spheroid partially expressing Lifeact-GFP in its ectoderm, introduced into the channel in a perpendicular configuration 3 hours after excision (Fig. 1C). Overlaid brightfield and spinning-disk projected images of the Lifeact-GFP signal (green) are shown. The tissue reorients its body axis and primary nematic fiber orientation along the channel axis and regenerates into a *Hydra* with a normal morphology. The elapsed time from excision is displayed (hh:mm), and the scale bar is 100 µm.

**Movie 4: Chimeric *Hydra* spheroid confined in a perpendicular configuration regenerating into a multi-headed morphology.** Time-lapse, spinning-disk confocal movie of a chimeric tissue spheroid partially expressing Lifeact-GFP in its ectoderm, introduced into the channel in a perpendicular configuration 3.5 hours after excision. Overlaid brightfield and spinning-disk projected images of the Lifeact-GFP signal (green) are shown. The tissue reorients its primary nematic fiber orientation along the channel axis and regenerates into a *Hydra* with two heads. The elapsed time from excision is displayed (hh:mm), and the scale bar is 100 µm.

**Movie 5: Regenerating chimeric *Hydra* spheroid confined in a perpendicular configuration transitioning from stable confinement to erratic movement within the channel.** Time-lapse, spinning-disk confocal movie of a chimeric tissue spheroid, partially expressing Lifeact-GFP in its ectoderm, introduced into a channel in a perpendicular configuration 3.5 hours after excision (Fig. S4). Overlaid brightfield and spinning-disk projected images of the Lifeact-GFP signal (green) are shown. The tissue remains stable for ∼24h, during which it reorients its primary nematic fiber orientation along the channel axis. Subsequently, the sample undergoes an abrupt transition and starts rotating erratically in the channel. The elapsed time from excision is displayed (hh:mm), and the scale bar is 100 µm.

**Movie 6: Reorientation of the actomyosin fibers in a confined regenerating *Hydra*.** Time-lapse, spinning-disk confocal movie of a regenerating *Hydra* expressing Lifeact-GFP in the ectoderm, introduced into a channel in a perpendicular configuration 3 hours after excision (Figs. 2-4). Top: Projected images of the basal surface of the ectoderm showing fiber organization in a regenerating *Hydra* spheroid confined in a perpendicular configuration. Bottom: Zoomed views of the boxed region showing projected views of the apical surface (left) and the basal surface (right). The fibers in the basal surface are color-coded according to their local orientation as in Fig. 3D. The tracked cells from the apical surface are depicted (dots). Tracking is discontinued at later time points because extensive tissue movements made reliable identification of individual cells difficult. The elapsed time from excision is displayed (hh:mm), and the scale bar is 100 µm.

**Movie 7: Actomyosin fiber reorientation in a confined chimeric *Hydra* spheroid.** Time-lapse, spinning-disk confocal movie of a chimeric *Hydra* partially expressing Lifeact-GFP in its ectoderm, introduced into a channel in a perpendicular configuration 4 hours after excision (Fig. 5). Top: Brightfield and fluorescence overlaid images of the fiber organization in the confined chimeric sample. Bottom: Zoomed projected images of the boxed region, which is centered around the tracked cells. Left: images of the apical surface of the ectoderm. Colored dots depict the position of manually tracked individual cells and the red line depicts the graft seam in the apical surface. Right: images of the basal surface of the ectoderm highlighting the fiber reorganization. The tracked cells and graft seam in the apical surface are indicated. Fibers in the protruding regions, defined as regions beyond the red line, are color coded according to their local orientation, as in Fig. 5C, highlighting the reorientation process. The elapsed time from excision is displayed (hh:mm), and the scale bar is 100 µm.

## Supplementary Note

We describe a minimal model for a nematic field embedded in an elastic sheet subject to an externally imposed uniaxial strain, incorporating both Frank elasticity and an alignment of the nematic field orientation to the elastic strain of the sheet. We use this model to compare the patterns of the nematic field governed by Frank elasticity alone with the behavior obtained when strain-alignment is also included.

### Model definition

We consider a two-dimensional linearly-elastic sheet with embedded fibers whose orientation is described by a nematic director

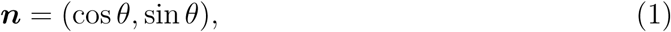

where *θ* is the fiber orientation angle. The nematic order parameter is defined at each point in space as

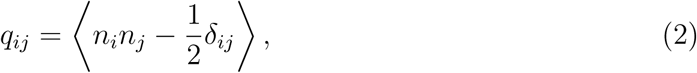

representing the coarse-grained fiber organization.

The deformation field of the elastic sheet is characterized by a displacement gradient tensor *∂_i_u_j_*, where *u_i_* is the displacement field. The pure shear strain is defined as the symmetric traceless part of the displacement gradient tensor

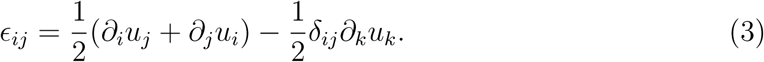

We consider the following setup:

- For simplicity, we assume that the nematic and strain fields are translationally invariant along the *x*-axis, so that all derivatives in the *x*-direction vanish.
- The elastic sheet is subject to a constant imposed pure shear strain *ɛ_xx_ >* 0, emu-lating the effect of geometric confinement on an elastic tissue spheroid in a channel.
- The nematic orientation is anchored perpendicular to the *x*-axis at the boundary at *y* = *L*, representing the orientation of the inherited fibers in the perpendicular configuration, where the inherited fibers are initially orthogonal to the channel. We choose the boundary condition for the nematic orientation at the opposite boundary at *y* = 0 to be parallel to the *x*-axis. This frustrated setting allows us to study how the competition between strain alignment and nematic Frank elasticity can lead to the formation of a domain boundary.

### Nematic field with Frank elasticity

The Frank energy of a nematic field describes the energetic cost of spatial variations of the nematic orientation. The corresponding effective stiffness, denoted Frank elasticity, arises from the microscopic interactions between the fibers and their tendency to align with each other, see Ref. 31 of the main text. The dynamical equation for the nematic field describing its relaxation towards equilibrium, considering only the Frank elasticity in the one-constant approximation, is given by

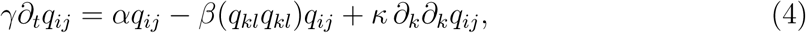

where *γ* is a rotational friction, *α >* 0 and *β >* 0 set the local nematic ordering, and *κ* is the Frank elastic constant. Furthermore, we consider a nematic that is deep in the ordered phase and approximate its norm to be constant, 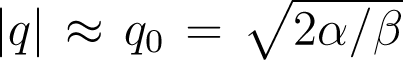. The dynamical equation in terms of the nematic angle *θ* is then

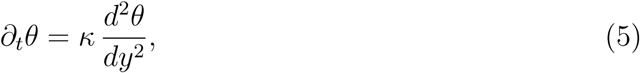

with the stable steady state solution

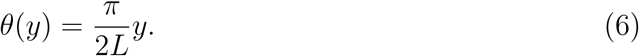

Therefore, Frank elasticity alone leads to a uniform gradient of the nematic orientation field across the entire system.

### Nematic field with strain-alignment and Frank elasticity

The dynamical equation for the nematic orientation angle, which includes both Frank elasticity and a strain-alignment term, is given by

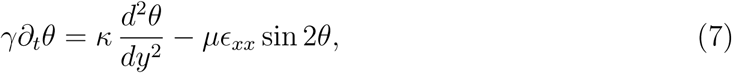

where we have introduced *µ* as the strain-alignment coefficient and *ɛ_xx_* as the imposed pure shear strain. At steady state, this equation becomes the well-known sine-Gordon equation describing the balance between Frank elasticity and strain-alignment interactions,

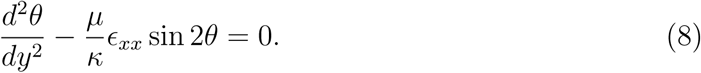

To analytically characterise the boundary layer forming at *y* = *L*, we approximate the system to be semi-infinite with

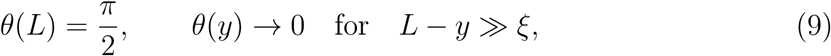

where *ξ* is the width of the boundary layer.

Multiplying the steady state equation by *dθ/dy* gives

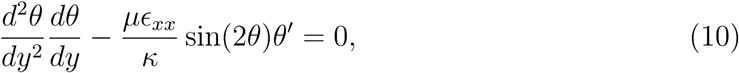

or equivalently

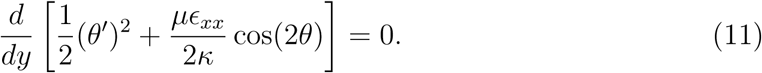

In the semi-infinite plane approximation, see Eq. 9, the boundary condition at *y* → −∞ corresponds to *θ*(−∞) → 0 and *dθ/dy*(−∞) → 0. This boundary condition determines the integration constant and yields

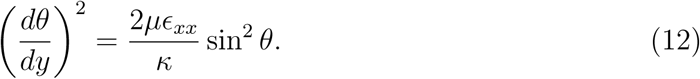

There are two solutions to this quadratic equation for *dθ/dy*, corresponding to the nematic orientation angle *θ* changing clockwise and counter-clockwise with increasing *y*−coordinate. Here, we consider the branch where the angle increases with increasing *y* coordinate

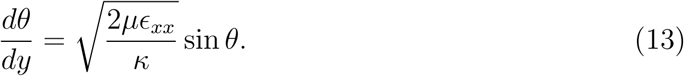

Separating variables,

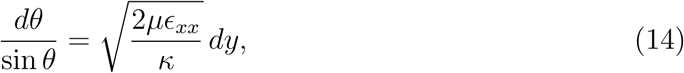

and integrating from the boundary at *L* to an arbitrary point *y* gives

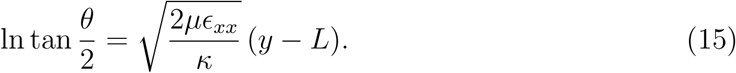

Therefore, the boundary-layer profile is

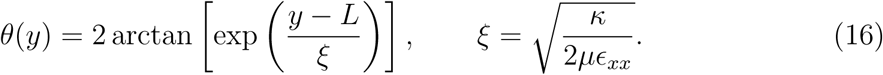

Thus the imposed strain introduces an intrinsic screening length *ξ*. Unlike the case with Frank elasticity alone, here the influence of the perpendicular boundary is localized near *y* = *L* when *L* ≫ *ξ*.

### Relevance for the nematic organization in confined ***Hydra*** spheroids

The actomyosin fibers that form in the initially disordered regions after a *Hydra* spheroid is inserted into a narrow channel are oriented along the channel axis. This orientation coincides with the direction of imposed tissue strain and is transverse to the adjacent inherited fibers, which are oriented perpendicular to the channel. As a result, narrow domain boundaries form between the newly formed parallel fiber arrays and the inherited, ordered transverse fiber domains.

This initial observed nematic pattern already indicates that the usual Frank elastic energy is not sufficient to describe the fiber organization. A narrow boundary between two perpendicularly oriented nematic domains is highly costly in terms of the nematic Frank energy. Furthermore, the orientation of the newly formed fibers along the tissue strain axis suggests the importance of a strain-dependent orientational coupling in the nematic model. The analysis of the simple model presented in this note illustrates that including strain-alignment interactions leads to the formation of a domain wall between two perpendicularly oriented domains with a well-defined wall thickness, *ξ*. This contrasts with a nematic governed by Frank elasticity only, in which the orientation field would turn gradually between the two orientations across the entire system.

## Notes

### Competing Interest Statement

The authors have declared no competing interest.

